# Single-Injection Multi-Omics Analysis by Direct Infusion Mass Spectrometry

**DOI:** 10.1101/2023.06.26.546628

**Authors:** Yuming Jiang, Ivan Salladay-Perez, Amanda Momenzadeh, Jesús Muñoz-Estrada, Uktarsh Tripathi, Anthony J. Covarrubias, Jesse G. Meyer

## Abstract

Combined multi-omics analysis of proteomics, metabolomics, and lipidomics requires separate liquid chromatography–mass spectrometry (LC–MS), which limits throughput and increases costs, hindering the application of mass spectrometry-based multi-omics to large-scale analyses. Here, we present <u>s</u>ingle-injection <u>m</u>ulti-omics <u>a</u>nalysis by <u>d</u>irect infusion (SMAD), an integrated platform leveraging ion mobility mass spectrometry and self-developed software tools to enable single injection multi-omics analysis without liquid chromatography. SMAD allows quantification of over 9,000 metabolite *m/z* features and over 1,300 proteins from the same sample in less than five minutes. We validated the efficiency and reliability of SMAD with three case studies. (1) mouse macrophages after M1/M2 polarization and senescence, (2) a pilot drug screen in human cells, and (3) large-scale high-throughput drug screening of mammalian cells in 96-well plates. Finally, relationships between proteomic and metabolomic data are discovered by machine learning and validated.

## Introduction

Multi-omics analysis and integration, which involves the analysis of at least a pair of genomic, epigenomic, transcriptomic, proteomic, lipidomic, and metabolomic data, has become increasingly essential for gaining a comprehensive understanding of biological processes and disease progression.^1, 2^ Currently, the combined use of LC and MS is the prevailing technology for proteome, metabolome and lipidome analysis.^3–5^ Separation of analytes with LC before MS is important to increase sensitivity and coverage, which enables detection of over 10,000 protein groups or thousands of metabolites from separate injection.^6–9^ However, despite advances in LC to enable shorter gradients, LC ultimately limits throughput of MS-based omics because of separation time followed by time for column washing and equilibration, and also due to requirements for different LC configurations for each omics layer.

The logical extreme of shorter LC is to remove LC completely and analyze molecules directly by direct infusion, which has already been demonstrated for both proteome and metabolome analysis.^10–12^ However, there exist two key challenges that restrict the coverage and depth of direct infusion mass spectrometry (DI-MS) methods: (1) ion suppression at the ion source caused by variations in ionization efficiency or abundance of ionizable analytes, and (2) ion competition effect in the mass analyzer, where high abundant ions conceal lower abundance ones.

Recent improvements in MS instrumentation (acquisition speed, mass resolution and sensitivity),^13–19^ advancements in mass spectrum interpretation software,^20–28^ and the integration of MS and ion mobility techniques (FAIMS, TIMS)^29–32^ have encouraged us to revisit the potential of DI-MS. For example, a spectral-stitching DI-MS method, which measures data as a series of mass-to-charge (*m/z*) intervals that are subsequently ‘stitched’ together to create a full mass spectrum, realized a total of ∼9,000 lipidome and metabolome *m/z* features in ∼5 min.^33^ An additional innovative technique for direct infusion metabolomic analysis involves using computational analysis to determine the optimal scan ranges that would yield the greatest number of *m/z* features.^34^ For direct infusion proteomic analysis, we originally described a rapid quantitative proteome analysis method by using two gas-phase separations by ion mobility and quadrupole selection, which identified over 500 proteins and quantified over 300 proteins in up to 3 min of acquisition time per sample.^35^ In a series of subsequent works, combined with our newly developed software CsoDIAq,^36^ we further improved the performance of this technology to more than 2,000 protein identifications, of which 1,100 were quantified.^37, 38^ There are also several previous instances of proteome analysis using direct infusion.^39–41^ However, the methods described above all focus on a single omics layer. They are insufficient to meet current demands for multi-omics analysis, specifically the rapid analysis of multiple omics components originating from the same sample simultaneously.^42–44^

Because proteomics, polar metabolomics, and lipidomics each require different LC configurations, the removal of LC presents an opportunity for combined analysis of all three omics layers. In this article, we describe a single-injection multi-omics analysis by direct infusion mass spectrometry (SMAD-MS) that integrates metabolome, lipidome, and proteome analysis in a single shot from the same sample. We replace LC with an extra gas-phase separation by high-field asymmetric waveform ion mobility spectrometry (FAIMS). We applied data-independent acquisition mass spectrometry (DIA-MS) for proteome analysis and spectral stitching of quadrupole slices with MS1 measurement for metabolome analysis, respectively. As we previously demonstrated for proteomics, we found that quadrupole slices combined with FAIMS separation enabled the detection of the most unique metabolite features. With this protocol, we achieved more than 1,300 protein identifications and detected over 9,000 metabolite m/z features from the same sample in just 5 minutes of total data acquisition. In two proof-of-principle applications across three case studies, we first demonstrate themacro profiling of multi-omics variation of macrophages after different polarizations or senescence, revealing significant multi-omics dysregulation and interaction. Secondly, in two additional case studies, we perform high-throughput screening of human cellular multi-omics responses to various drug treatments with 96-well plates, followed by further machine learning integration of metabolome and proteome changes. Our approach provides a straightforward, high-throughput, and cost-effective option for multi-omics analysis.

## Results

### Overview of Single-shot Multi-Omics approach by Direct Infusion Mass Spectrometry (SMAD-MS)

The principled schematic and workflow of the SMAD method are shown in **Fig. 1a**. In brief, different omics isolated from the same sample were mixed and then directly infused to the ion source without any separation in liquid chromatography. Following the ionization, two gas-phase separation techniques were utilized with the aim of reducing the complexity of gas-phase ions before they entered into the mass analyzer, namely, (1) FAIMS separates ions depending on the compensation voltage and (2) the quadrupole separates ions according to their mass-to-charge ratio (*m/z*). Ultimately, all gas-phase ions are sequentially detected by the orbitrap.

**Fig.1.**
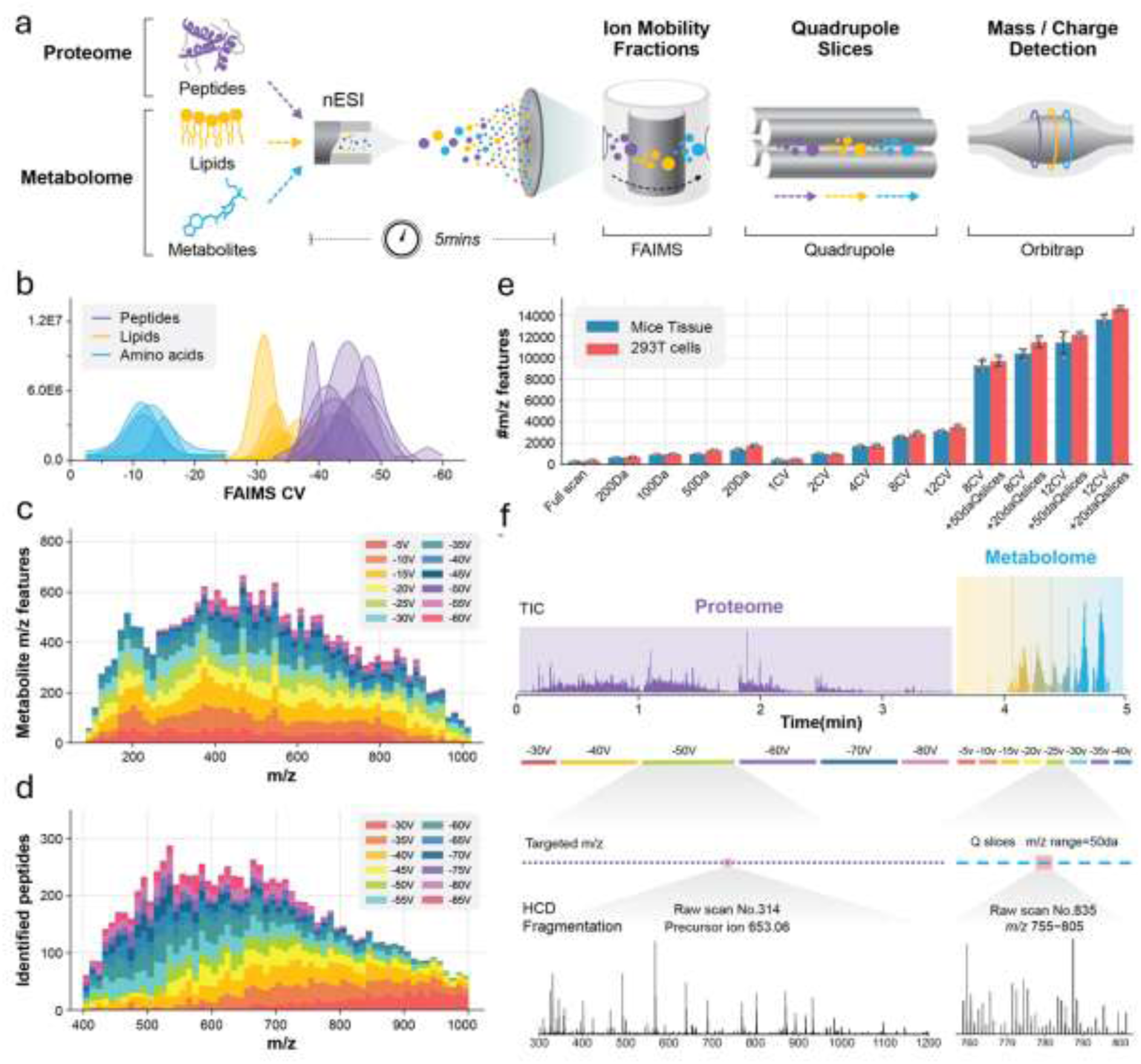
| Overview of SMAD-MS workflow. **a,** Scheme of our high-throughput approach for single-injection analysis of peptides and metabolites using SMAD-MS. **b,** Typical compensation voltage (CV) intervals of peptide, amino acid, and lipid standards in FAIMS CV space. The raw data traces over CV space were smoothed to approximate Gaussian distributions. **c, d**, Number of detected features in different *m/z* intervals and FAIMS compensation voltages (CV) from a real metabolome (c) and proteome (d) sample derived from 293T cells. **e,** Number of detected metabolite features under various compensation voltages and quadrupole slices of samples extracted from mouse tissue and 293T cells. Data are from three repeated injections of the same sample (n=1) with different parameters. **f,** Schematic diagram of typical experimental settings for SMAD, including a typical TIC diagram, CV settings, targeted m/z, Q slices, and typical mass spectrum of proteome and metabolome, respectively. The entire data collection process requires approximately 5 minutes per sample for acquisition time.

We first tested the compensation voltage (CV) range of different standard molecules and results demonstrated that lipids, amino acids, and peptides occupy different optimum CV intervals (**Fig. 1b**), for example, amino acids transmit through FAIMS in the range of −5V to −20V, lipids transmit in −20V to −35V, and peptides transmit from −30V to −70V. A more complexed real metabolome and proteome sample derived from 293T cells further proved that the optimum CV interval for metabolome (metabolites, lipids) and proteome (peptides) are 10-35V, and 40-60V, respectively (**Fig. 1c, d**). This result enables the fractionation of molecules originating from distinct omic sources, thereby minimizing interference between them.

Next, we assessed the impact of gas phase separation using FAIMS compensation voltages and/or quadrupole slices on the number of detected metabolite m/z features from samples extracted from mouse tissue and 293T cells, as we did previously for peptides from the proteome.^35^ The number of *m/z* features detected increased with the number of FAIMS CVs or with decreasing quadrupole isolation window width (**Fig 1e**). The use of both FAIMS and quadrupole slices further increases the number of unique detectable metabolite *m/z* features, which provided confirmation of significant ion competition in the Orbitrap and highlighted the necessity of multiple gas-phase separations before mass analysis.

Based on these findings, we established the final experimental settings of SMAD as depicted in **Fig. 1f**. We applied data-independent acquisition mass spectrometry (DIA-MS) for proteome analysis and spectral-stitching quadrupole slices with MS1 measurement for metabolome analysis. We utilized six FAIMS CVs ranging from −30V to −80V in a step of 10V for proteome acquisition, and −5V to −40V with a 5V step for metabolome analysis. It is worth noting that the instrument parameters, such as mass resolution, compensation voltages, and the number of target peptides, are adjustable and can be modified to match the specific requirements of the experiment and the sample type. Generally, the total acquisition time does not exceed five minutes per sample.

### Data processing, performance optimization and quantitative evaluation of SMAD

**Fig. 2a** summarizes our initial data processing workflow for raw files produced by SMAD. MS1 data from the metabolome part are extracted from the RAW file during transformation to mzML.^45^ The direct infusion data is formatted such that retention time corresponds to decreasing FAIMS compensation voltage, allowing extraction of extracted ion mobiligrams (XIMs) for quantification using MZmine3 (**Fig. S1**).^21^ MS/MS data from the proteome part is converted to mzXML for analysis using ZoDIAq software ^36, 37^ (https://github.com/xomicsdatascience/zoDIAq) to identify and quantify peptides and proteins. This data analysis workflow was used to assess the performance of SMAD.

**Fig.2.**
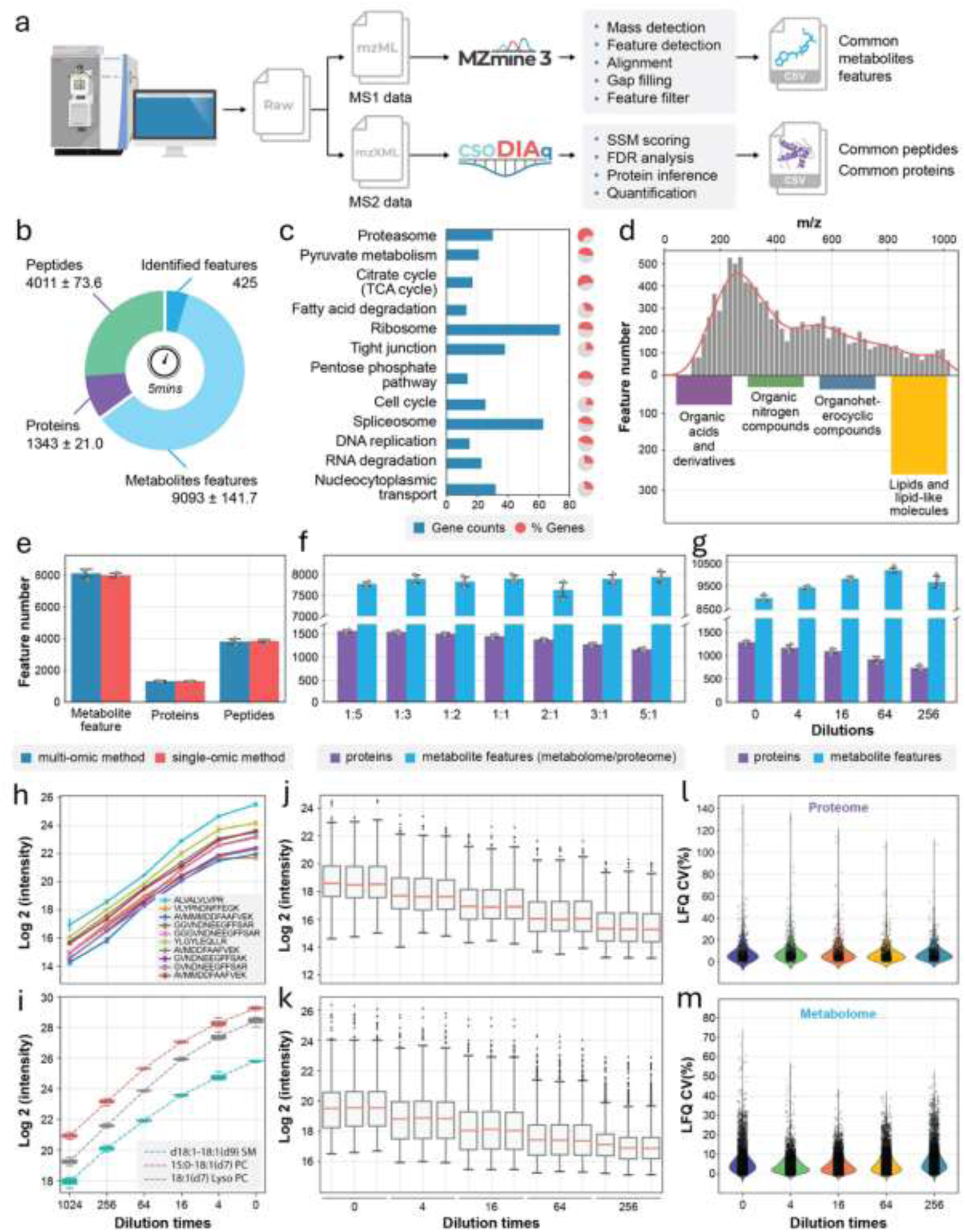
| Data processing, performance optimization, and quantitative evaluation of SMAD. **a,** Scheme showing data processing flow starting from a typical raw file produced by SMAD. **b,** Number of detected metabolite features, peptides, and proteins in a 5-minute acquisition time by SMAD of extracts from 293T cells. Counts are the average plus or minus the standard deviation from three independent biological samples (n=3). **c,** Proteins identified by SMAD across three replicate samples from 293T cells were analyzed by KEGG pathway enrichment analysis. The bars indicate the number of proteins identified in the pathway, and the colored proportion of the circle reflects the coverage of proteins in each pathway. **d,** *m/z* distribution of detected metabolite features and classes of identified metabolite features by SMAD of 293T cells. **e,** The comparison of detected molecule features (metabolites, peptides, proteins) between a single-injection multi-omics acquisition method and a separate single-omics acquisition method. **f,** The effect of different mixing proportions of peptides and metabolites on detected molecule features by SMAD. (metabolome/proteome) **g,** Performance of SMAD in detecting different concentrations. Original concentration is proteome samples dissolved in 40μl metabolome extraction (produced by adding 500μl solvent to 2million cells) and maintaining the final proteome concentration to 4μg/μl. For different methods, mixing ratios, and dilutions, data from each condition were collected from three repeated injections of the same sample (n = 1). **h,i,** label free quantification curve of standard peptides from MS-QCAL protein (**h**) and standard lipids from Avanta (**i**) at different concentrations. Data are from three repeated injections of the same sample (n=1) for each dilution. The value from each replicate was determined as the average of at least 20 scans of that molecule in that injection. **j,k** Untargeted quantification of detected proteins and metabolite features was performed using SMAD (each concentration was measured in technical triplicate, n = 1). **l,m,** coefficient of variation of all quantified proteins and metabolite features for different dilutions.

SMAD collected in only five minutes per 293T sample produced an average of 4,011 peptides, 1,343 protein groups, and 9,093 metabolite *m/z* features (out of which 425 were identified) (**Fig. 2b**). For the identified protein groups, Kyoto Encyclopedia of Genes and Genomes (KEGG) pathway enrichment analysis revealed numerous important cellular pathways, including central carbon metabolism (i.e., tricarboxylic acid cycle, pyruvate metabolism, pentose phosphate pathway), protein synthesis and degradation (i.e., ribosome, spliceosome and proteasome), and nucleic acid replication and transport (DNA replication, RNA degradation and nucleocytoplasmic transport). For TCA cycle and proteasome pathways, the identified proteins accounted for more than 50% of all related genes in those specific pathways (**Fig. 2c and Fig. S2**). The metabolite feature *m/z* distribution shows that most signals are within the interval from 200 to 400 *m/z* (**Fig. 2d**). We separately performed MS/MS of same sample attempting to identify some metabolite features quantified by SMAD. Results demonstrated that many lipids, as well as organic acids and organic nitrogen compounds, were identified (**Fig. 2d**). We also applied this method to other sample types. Similar to the results obtained from 293T cells, the analysis of macrophages yielded a comparable depth of proteome coverage (over 1,300 proteins) and metabolite m/z features (∼9,000), as well as a broad representation of cellular pathways (**Fig. S3**).

We also wondered whether SMAD would result in a significant reduction in the number of detected molecule features compared to separately analyzing the metabolome and proteome fractions by direct infusion. The results show that the number of detected peptides, proteins, and metabolite features is essentially the same level between the two different methods (**Fig. 2e**), as well as protein species and *m/z* distribution of metabolite features (**Fig. S4a-c**). Additionally, we investigated whether variations in the mixing ratio of two different omics samples could affect the performance of SMAD. Notably, we examined seven samples with various mixing ratios across a 25-fold (metabolome/proteome volume ratio). We observed that, as the proportion of metabolome increased, the number of identified proteins decreased by up to 30%. However, the quantity of detected metabolite features remained relatively constant (**Fig. 2f**). Furthermore, we also investigated the influence of the total sample concentration. The results show that as the concentration of the sample decreases, the number of detected proteins also decreases, but the detected metabolites show a trend of increasing first and then decreasing (**Fig. 2g**). These results indicate that protein identifications, namely, the proteome part, is more sensitive to sample mixing ratios and concentrations. The potential reason is that the signal intensity of tandem mass spectrometry in proteomic analysis is much lower than that of the precursor signal of metabolites, resulting in a more sensitive phenomenon to factors such as concentration. The influences of these factors for protein groups, protein functions, and metabolite *m/z* distribution were illustrated in **Fig. S4d-i.**

The direct infusion strategy completely omitted liquid chromatography, resulting in the absence of retention times or elution profiles that could be used for peak integration. Here, we implemented a label-free quantitative strategy for both the metabolome and proteome, based on XIM area and peptide fragments, respectively (**Fig. S1**). Starting with a mixture of 3 lipids and 10 QCAL peptide standards, the results showed an excellent linearity across a broad range of concentrations (**Fig 2h, i**). We further spiked these mixture standards into real multi-omics sample derived from HEK293T cells and surprisingly, the standards remain good linearity across different concentrations (**Fig. S5a-d**). Furthermore, we applied the same strategy for the whole proteome and metabolome, quantification results exhibit excellent repeatability (**Fig. S5e, f**) and linearity across all 437 proteins and 3970 metabolites *m/z* features (**Fig. 2j, k**) and typical proteins and metabolites were also shown in **Fig. S5g, h.** The coefficient of variance analysis at different concentrations demonstrated that 90.6% proteins and 93.4% metabolite *m/z* features had CV less than 0.2, which further proved the reliability and robustness of our LFQ strategy of SMAD (**Fig 2l, m**).

### Case Study 1: Accelerated Multi-Omics Profiling of Macrophage Polarization and Irradiation Responses via the SMAD Platform

To demonstrate the potential of SMAD for real biological samples, here we conducted the first case study focused on the multi-omics responses of macrophages following different polarization and irradiation. Macrophages are well known to differentiate into either the pro-inflammatory M1 or anti-inflammatory M2 polarization states after treatment with lipopolysaccharide (LPS) and interleukin-4 (IL-4), respectively.^46, 47^ We started with this system as a positive control where we should be able to verify results with literature context. Beyond these established treatments, irradiation was used to generate a more novel senescent phenotype. We applied SMAD to understand the molecular mechanisms underlying these changes. Mouse bone-marrow-derived macrophages (BMDMs) were treated with immune stimulators LPS and IL-4 or irradiated in 10 cm dishes for 24 hours. After washing with PBS, lipids and metabolites were extracted by a mixed solvent (ISO/ACN/H2O, 4:4:2, volume ratio), then the precipitated protein pellet from the same sample was further processed (lysis, digest and desalt) for proteomics into peptides (**Fig. 3a**). We measured the multi-omics by SMAD. A total of 1,386 protein groups were identified. We also detected 9,829 metabolite *m/z* features, and 541 of them were identified using GNPS^22^ (**Fig. 3b**, **Fig. S6, Supplementary Table 1, See methods**). Coefficient of variation (CV) analysis demonstrated the robustness and reproducibility of the SMAD method, with median CVs of 0.18 for quantified proteins and 0.21 for metabolite *m/z* features (**Fig. 3c**).

**Fig.3.**
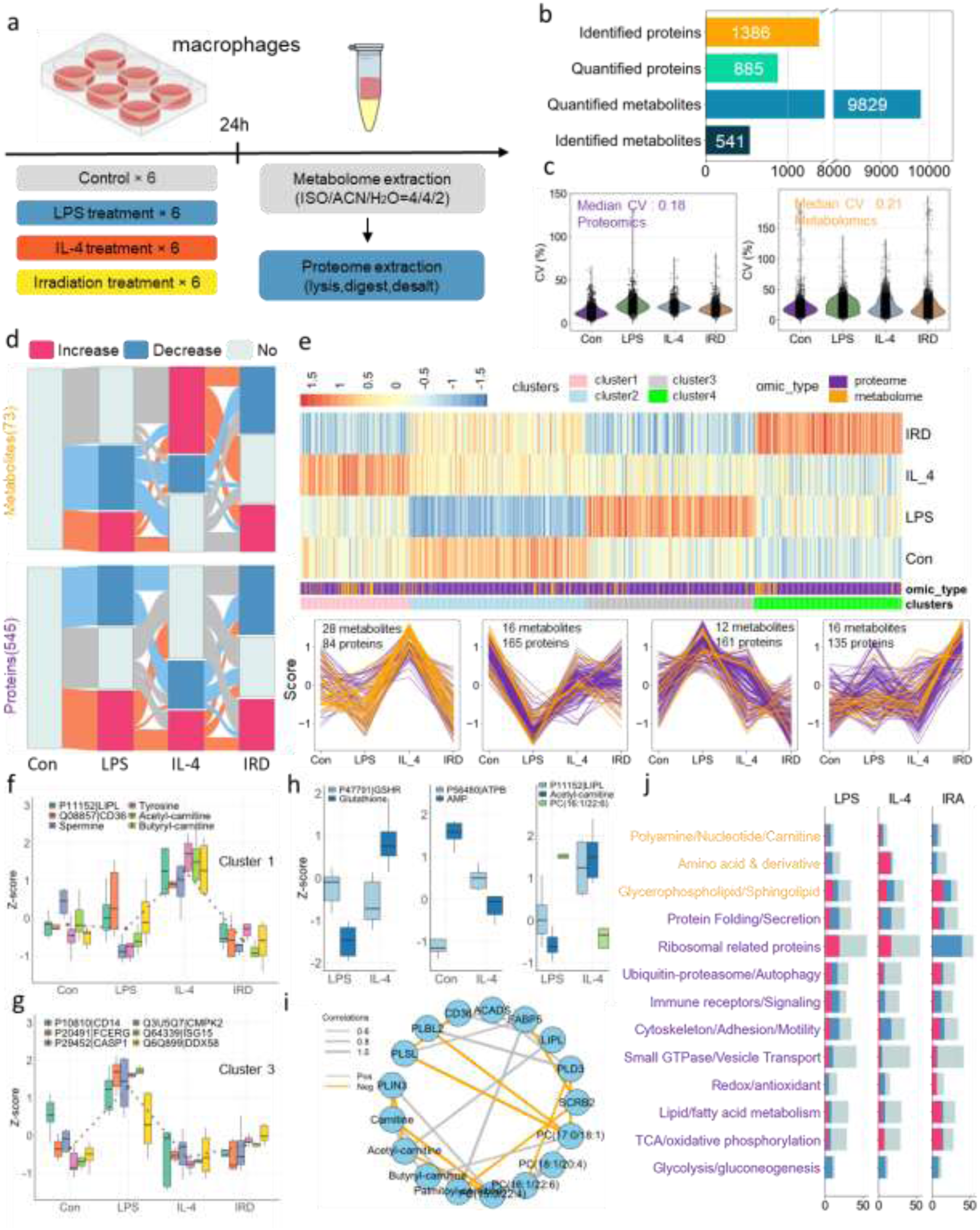
| SMAD enabled rapid multi-omics analysis of macrophage activation and radiation-induced perturbations. **a**, Schematic diagram of experimental design. (LPS: lipopolysaccharide, IL-4: Interleukin 4, IRA: irradiation). **b**, Overview of molecules monitored by SMAD. **c**, Coefficient of variance distribution for all identified proteins and metabolite features across all samples within each treatment. **d**, Changes of significant dysregulated molecules (metabolites and proteins) among three treatments and control. (two-sided Wilcoxon rank test with Benjamini–Hochberg (BH) adjusted P values <0.05). **e**, The Heatmap and clustering of dysregulated multi-omics molecules following immune activation or irradiation. (K-means clustering). **f,g,** Typical dysregulated multi-omics molecules in cluster1(f) and cluster3(g). **h,** Typical dysregulated multi-omics feature pairs after polarization/irradiation. The box shows the quartiles of the dataset while the whiskers extend to show the rest of the distribution. (n=6 independent biological samples for each treatment and control for a total of 24 biological samples). **i,** Pearson correlation network analysis of multi-omics molecule features involved in lipid metabolism. Each node represents a molecule feature, and the edge width represents the correlation strength. **j,** Stacked overview of boxplots showing the perturbation patterns of multi-omics pathways and metabolic molecules in macrophages after polarization or irradiation.

Before performing further multi-omics data analyses, we aimed to verify the reliability of our newly developed SMAD method. To achieve this, we analyzed the same set of macrophage samples using a conventional LC-MS proteomics approach (**Supplementary Table 2**). Dimensionality reduction analysis of the proteomics datasets obtained from both methods revealed highly similar distribution patterns among treatment groups (**Fig. S7a, b**). Furthermore, quantitative comparisons of individual protein levels, as well as the ranking patterns of protein intensities across treatment groups, showed high consistency between the two methods, with correlation coefficients exceeding 0.7 (**Fig. S7c-e**). Additionally, Pearson correlation analysis demonstrated that over 80% of the identified proteins exhibited consistent patterns of variation across different treatment conditions (**Fig. S7f-i**). Collectively, these findings indicate that the results obtained from the SMAD method closely align with those from conventional LC-MS analysis, thereby further substantiating the reliability of our newly developed methodology.

Then, identified proteins and metabolites from SMAD were first analyzed using one-way ANOVA, revealing that 545 proteins and 73 metabolites exhibited significant alterations with either treatment (Benjamini–Hochberg (BH)-adjusted P values <0.05) (**Fig. 3d**). KEGG pathway enrichment demonstrated that the significant changed proteins are related to essential pathways including glycolysis, TCA cycle, oxidative phosphorylation, Cholesterol metabolism, mitophagy, spliceosome and endocytosis. (**Fig. S8a**). Further two-sided Wilcoxon rank-sum tests between each condition and control revealed that although some molecular features exhibited similar alterations, a larger proportion displayed divergent or even opposite dysregulation patterns in response to different treatments (**Fig. 3d**). Complementary PCA and correlation analyses of the multi-omics data also clearly distinguished the treatments from one another and demonstrated low correlations for molecules among treatments (**Extended Data Fig. S8b, c**). These findings highlight distinct multi-omic characteristics of macrophages after different polarization states or irradiation, aligning well with established knowledge.^48^

Next, to better interpret the variations in metabolites and proteins related to immune activation and irradiation, we utilized K-means clustering to analyze the identified significant molecule features from the ANOVA. The significantly dysregulated molecules were categorized into four primary clusters (**Fig. 3e**). Specifically, cluster 1 primarily consisted of molecules (84 proteins and 28 metabolites) that were upregulated in response to IL-4 (**Fig. 3e**). These features were associated with significant pathways, including energy metabolism, immune response receptors, and fatty acid metabolism. For example, several amino acids (histidine, lysine, tyrosine, arginine), carnitine derivatives (acetyl-carnitine, butyryl-carnitine), and lipid metabolism–related proteins (CD36, LIPL) were significantly upregulated (**Fig. 3f, Fig. S8d**). The increased levels of amino acids may support biosynthetic and immunoregulatory processes involved in tissue repair. Elevated carnitine derivatives, along with CD36 and LIPL, suggest enhanced fatty acid uptake and β-oxidation, contributing to sustained energy production through oxidative phosphorylation. Together, these changes reflect a metabolic shift toward anabolic and oxidative metabolism, characteristic of M2 macrophages engaged in anti-inflammatory responses and tissue remodeling.

In addition, cluster 3, which included 12 metabolites and 165 proteins, exhibited a significant increase in response to LPS treatment, indicating a significant upregulation following M1 polarization. The proteome features are mainly related to KEGG pathways such as protein processing in endoplasmic reticulum, endocytosis and spliceosome (**Fig. S8e**). Several immune-related proteins were consistently upregulated (**Fig. 3g**), including CD14 (TLR4 co-receptor), DDX58/RIG-I (viral RNA sensor), ISG15 (ubiquitin-like inflammatory regulator), and CMPK2 (mitochondrial kinase linked to nucleotide metabolism and immune activation). FCERG (immunoreceptor signaling) and CASP1 (IL-1β–processing inflammasome component) were also elevated, reflecting strong activation of innate immune pathways, consistent with previous reports of M1 macrophage polarization. ^49^

In contrast, Cluster 2 contained 16 metabolites and 165 proteins significantly downregulated under M1 polarization. Reductions in glutathione and carnitine suggest a decrease in antioxidant capacity and fatty acid oxidation, in line with the glycolytic shift of pro-inflammatory macrophages (**Fig. S8f**). AMP levels were also reduced, indicating altered energy homeostasis and possible AMPK suppression. Protein-level changes included downregulation of ALDOA (a key glycolytic enzyme), 14-3-3 gamma (1433G), and beta-tubulin 4B (TBB4B), implicating disruptions in glycolysis, signaling, and cytoskeletal structure **(Fig. S8f, g)**. These coordinated molecular changes likely reflect comprehensive shifts in metabolic priorities, cellular signaling, and structural organization that accompany macrophage polarization toward an M1 inflammatory state.

The final category, Cluster 4, comprising 135 proteins and 16 metabolites, exhibited significant upregulation following irradiation. The increased expression of ATP synthase subunits (ATPK, ATPA, ATPB, ATP5H) suggests enhanced mitochondrial oxidative phosphorylation, potentially reflecting elevated energy demands associated with stress adaptation and cellular repair (**Fig. S8h**). Notably, the upregulation of β-galactosidase (BGAL), a lysosomal hydrolase involved in the degradation of glycosylated substrates, may indicate activation of autophagic flux. As a marker of lysosomal activity and a canonical marker of senescence, elevated BGAL expression supports the notion of enhanced senescence and autophagy, which may facilitate the clearance of damaged cellular components and contribute to metabolic remodeling in irradiated macrophages (**Fig. S8h, i, j**).

Furthermore, we also observed several compelling instances of multi-omics co-perturbations. Notably, we identified coordinated dysregulations involving the redox-related protein glutathione reductase (GSHR) and the small molecule glutamine; the energy metabolism-related protein ATP synthase subunit beta (ATPB) paired with AMP; and the lipid metabolism-related protein lipoprotein lipase (LIPL), together with the small molecule carnitine and its associated lipid species (**Fig. 3h**). These co-perturbation patterns, spanning distinct omics layers, align closely with established biological insights. Building upon these observations, we specifically extracted multi-omics features related to lipid metabolism for further analysis. This analysis uncovered significant correlations among proteins involved in lipid metabolism, small molecules such as carnitine and its derivatives, and corresponding lipid species. For example, these proteins and carnitine derivatives, known for their collaborative roles in lipid catabolism, displayed strong positive correlations. Conversely, the associated lipid species showed negative correlations with these molecules, indicative of their depletion during lipid breakdown processes (**Fig. 3i, Fig. S9a**).

To better elucidate the dysregulation patterns of macrophages following M1/M2 polarization or irradiation, we further performed correlation network analysis of the identified proteins and metabolites (**Fig. S9b**), and classified them into distinct pathways, comprising ten proteomic pathways and three metabolomic pathways (**Fig. 3j**). The results revealed that proteins involved in the same pathway exhibited strong correlations, yet also revealed complex regulatory dynamics, with both upregulated and downregulated features observed within most multi-omics pathways. Interestingly, glycolysis-related proteins consistently exhibited downregulation across all three treatment conditions, a pattern that was also validated by the LC-MS results (**Fig. S10)**. This suggests that the general suppression of glycolytic metabolism may be associated with metabolic reprogramming during macrophage activation or stress responses (**Fig. 3j, Fig. S9c-e**). Notably, although classically activated M1 macrophages are typically characterized by a firm reliance on glycolysis, we observed a downregulation of glycolytic pathways even under M1 polarization. This unexpected pattern may reflect the specific time point of polarization or experimental conditions used, which may not fully sustain glycolytic activation. Alternatively, a negative feedback mechanism may have been engaged to limit glycolytic flux and avoid excessive inflammatory responses. It is also possible that macrophages undergo temporal metabolic transitions during polarization, initially upregulating glycolysis, followed by a later-phase attenuation, resulting in an overall downregulated profile at the measured time point. These findings highlight the importance of considering temporal dynamics and stimulation intensity when studying macrophage metabolic states.

Additionally, ribosomal proteins were specifically and significantly downregulated following irradiation (**Fig. 3j, Fig. S9f-h**), suggesting compromised protein synthesis under irradiation-induced cellular stress. Additionally, irradiation uniquely resulted in increased levels of redox-related proteins, reflecting an elevated oxidative stress response, whereas both M1 and M2 polarization showed an overall opposite trend, indicating divergent responses to oxidative balance (**Fig. 3j**). Moreover, several immune signaling and receptor-related proteins showed differential dysregulation patterns depending on the polarization state yet maintained strong correlations among functionally similar receptors (**Fig. S9i-k**). A marked upregulation of amino acids and their derivatives was also observed in the metabolomics data, specifically under M2 polarization (**Fig. 3j)**. This metabolic signature may reflect enhanced anabolic activity and biosynthetic readiness in M2 polarized macrophages, which are known to support tissue remodeling and repair. The increased availability of amino acids may also contribute to immunoregulatory functions, further distinguishing M2 macrophages from their pro-inflammatory M1 counterparts. Collectively, these findings underscore the significant metabolic flexibility and distinct reprogramming that macrophages undergo in response to polarization stimuli and irradiation stress. These views of this established system would not be possible by measuring a single omic layer, highlighting the value of this approach. Future research should focus on exploring the functional consequences of these metabolic alterations, particularly their implications in macrophage-driven inflammation, immunity, and tissue repair processes.

### Case Study 2: Drug-Induced Multi-Omics Responses in Human Cells

Rapid and high-throughput profiling of multi-omics responses to pharmacological perturbations is crucial in drug discovery and chemical biology. Leveraging the high-throughput capability of our SMAD platform, we further adapted it for systematic drug screening in 96-well plates. In this second case study, we applied SMAD to monitor the multi-omics responses of human 293T cells exposed to five distinct compounds: deferoxamine (an iron chelator), Torin2 (an mTOR inhibitor), ISRIB (an integrated stress response inhibitor), MG132 (a proteasome inhibitor), and A939572 (an SCD1 inhibitor). Cells were cultured in a 96-well plate and processed using a SMAD protocol optimized for high-throughput sample handling (**Fig. 4a**). The total MS acquisition time for one full plate was approximately 7 hours. SMAD enabled the identification of 1,616 proteins, with 838 proteins retained after data filtering (**Fig. S11a**). Additionally, a total of 7,005 metabolite features were quantified, among which 425 metabolites were confidently identified (**Fig. 4b, Fig. S11b-c, Supplementary Table 3**). Notably, over 80% of the quantified features exhibited a coefficient of variation (CV) below 0.5, with median CVs of 0.21 for proteomics and 0.20 for metabolomics, respectively (**Fig. 4c**). These results highlight the robustness and reproducibility of SMAD for high-throughput multi-omics drug screening.

**Fig.4.**
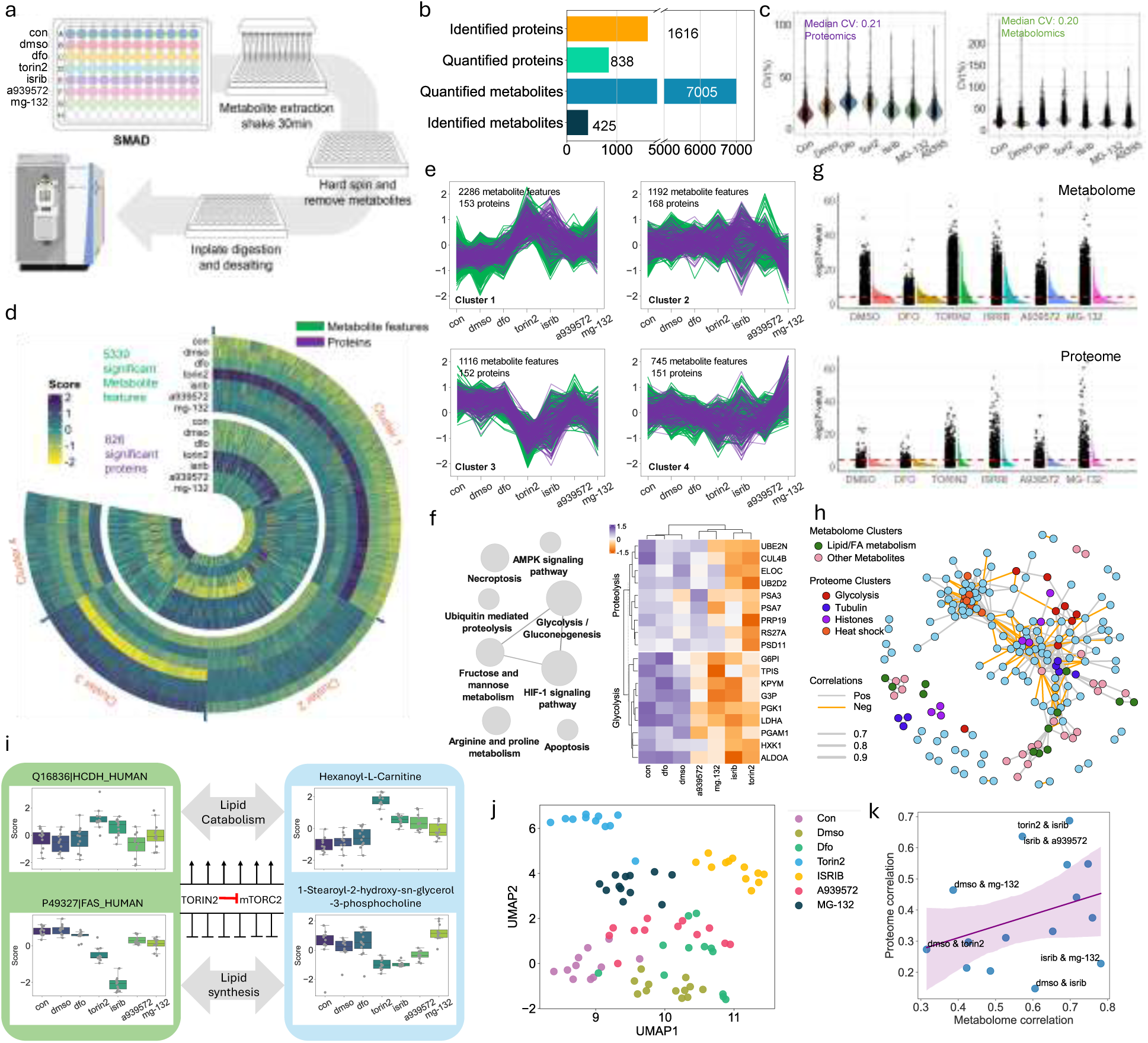
| High-Throughput SMAD-Based Profiling of Drug-Induced Multi-Omics Responses in Human Cells. **a,** Overview of experimental design and sample processing flow in 96-well plates. **b,** Molecular information and data quality of multi-omics dataset acquired by SMAD. including identified, quantified features. **c,** Coefficient of variance distribution for all identified proteins and metabolite features across all samples within each treatment. **d,** The Heatmap showing significant dysregulated multi-omics molecules among drug treatments (one-way ANOVA test with Benjamini–Hochberg (BH) adjusted P values <0.05). **e,** Four clusters of significantly dysregulated multi-omics molecules of fig4.d (K-means clustering). **f,** Enriched KEGG pathways(left) of cluster 3 and typical proteins in glycolysis and proteolysis (right). **g,** Raincloud of significantly dysregulated metabolome m/z features (up) and proteins (down) in different treatments. **h,** Correlation network of significantly dysregulated and identified multi-omics molecules (proteins and metabolites). Only Pearson correlations with an absolute value greater than 0.7 are included in this network, and typical clusters are labeled with a distinct color. **i,** A typical example of interaction between two omics layers: the regulation of lipid synthesis and catabolism after Torin2 treatment. The box shows the quartiles of the dataset while the whiskers extend to show the rest of the distribution. Data points are shown as dots. **j,** UMAP analysis of multi-omics responses among all drugs and controls. (All metabolite features and proteins included). **k,** The correlation between metabolic and proteomic levels across different drug pairs. All data in this figure comes from 82 independent samples corresponding to unique wells in the 96 well plate. n=11-12 per condition.

One-way ANOVA analysis was first applied to determine dysregulated proteins and metabolites after various drug treatments (Benjamini–Hochberg (BH)-adjusted P values <0.05). As shown in **Fig. 4d**, a total of 5339 metabolite features and 626 proteins were significantly altered in at least one treatment. To better illustrate the variations across various treatments, we consolidated all significantly dysregulated molecules and categorized them into four clusters utilizing K-means clustering (**Fig. 4e**). Cluster 1, which represents the profiles of 153 proteins and 2286 metabolite *m/z* features, generally peaked with TORIN2 or ISRIB treatment. Further analysis revealed that the disrupted proteins in this cluster are primarily associated with the pentose phosphate pathway, cell cycle, spliceosome, fatty acid degradation, and protein processing in the endoplasmic reticulum (**Fig. S11d,e**). This may be due to the repair of some protein synthesis processes and the increase in cellular energy consumption after ISRIB treatment. Additionally, cluster 3, enriched by 152 proteins and 1,116 metabolite *m/z* features, demonstrated an opposite trend that was downregulated at either of these two treatments. KEGG enrichment analysis of proteome results revealed that the dysregulated proteins are mainly related to pathways like glycolysis, HIF-1 signaling pathway, amino acid metabolism and ribosome (**Fig. 4f**), which can be attributed to the inhibition of cell growth by TORIN2. Cluster 4, which represents the profiles of 151 proteins and 745 metabolite *m/z* features, generally peaked at MG-132 treatment. MG132 is a proteasome inhibitor that blocks the activity of the proteasome. Here, we demonstrated that proteins associated with the cytoskeleton, spliceosome, and DNA replication were significantly upregulated in MG-132 (**Fig. S11f, g**). Moreover, Cluster 2 exhibited a relatively heterogeneous alteration pattern; however, it generally demonstrated a downregulation under MG-132 treatment and dysregulation in response to ISRIB (**Fig. S11h, i**).

Further analysis with t-tests compared to the control also revealed TORIN2, ISRIB, and MG-132 induced more changes to both the proteome and metabolome layer (**Fig. 4g**), while limited variations were observed between the control, DMSO, and DFO treatment. Consistent with our earlier findings, treatment with 10 µM DFO resulted in minimal alterations across multi-omics profiles.^37^ Correlation analysis of significantly altered molecules revealed that those involved in the same pathways or with similar functions tend to cluster together (**Fig. 4h**). This observation is consistent with the logic of cellular functional responses and further supports the effectiveness and accuracy of our method. Additionally, molecules from different omics layers also exhibited correlations, suggesting potential multi-omics connections and interactions (**Fig. 4h**). Furthermore, a multi-omics co-regulation related to fatty acid and lipid metabolism was discovered in the TORIN2 treatment group. We observed a significant upregulation of mitochondrial fatty acid beta-oxidation enzyme (Q16836, HADH) and, correspondingly, increased carnitine, acetyl-carnitine, and hexanoyl-carnitine enhanced the lipid catabolism (**Fig. 4i, Fig. S11j**). We also observed a notable decrease of fatty acid synthetase (P49327, FAS) in TORIN2 treated cells and typically lipids were accordingly downregulated in this treatment, which represents the inhibited lipid synthesis process (**Fig. 4i**). Notably, TORIN2 treatment was associated with molecular changes suggestive of decreased lipid synthesis but not decreased lipid catabolism.

UMAP dimensionality reduction of the multi-omics data from individual replicates revealed clear separation among treatment and control groups, highlighting distinct multi-omics profiles associated with each condition (**Fig. 4j**). To further investigate the cellular responses to different drug treatments, we assessed the correlations between metabolic and proteomic profiles across drug pairs. The analysis revealed that TORIN2 and ISRIB exhibited the highest correlation in proteomic alterations (Pearson correlation = 0.701), while TORIN2 and MG-132 showed the strongest correlation in metabolic changes (Pearson correlation = 0.729) (**Fig. 4k, Fig. S11k,l**). Notably, a relatively positive linear correlation was observed between metabolomic and proteomic responses, suggesting a potential intrinsic link and coordinated regulation between these two molecular layers.

### Case Study 3: SMAD enables integrative multi-omics strategies for large-scale drug screening

Encouraged by the successful application of the SMAD method demonstrated in Case Study 2, here we expanded this approach into a broader drug screening investigation to further underscore its practical utility and scalability. Specifically, we designed an experimental screen consisting of 72 distinct small-molecule compounds (**Fig. S12a**). To enhance the statistical robustness of the experimental results, we employed six replicates for each drug. Increasing the number of replicates significantly reduces the standard error, improves the power of statistical tests, and enhances the ability to detect true biological differences, thereby minimizing biases caused by random variability (**Fig. S13**). Additionally, up to 12 positive and negative controls were systematically incorporated into each 96-well plate to validate data quality and facilitate correction for potential batch effects (**Fig. 5a**). Following the sample processing workflow established in Case Study 2, metabolites and proteins were separately extracted and subsequently analyzed via mass spectrometry-based profiling.

**Fig.5.**
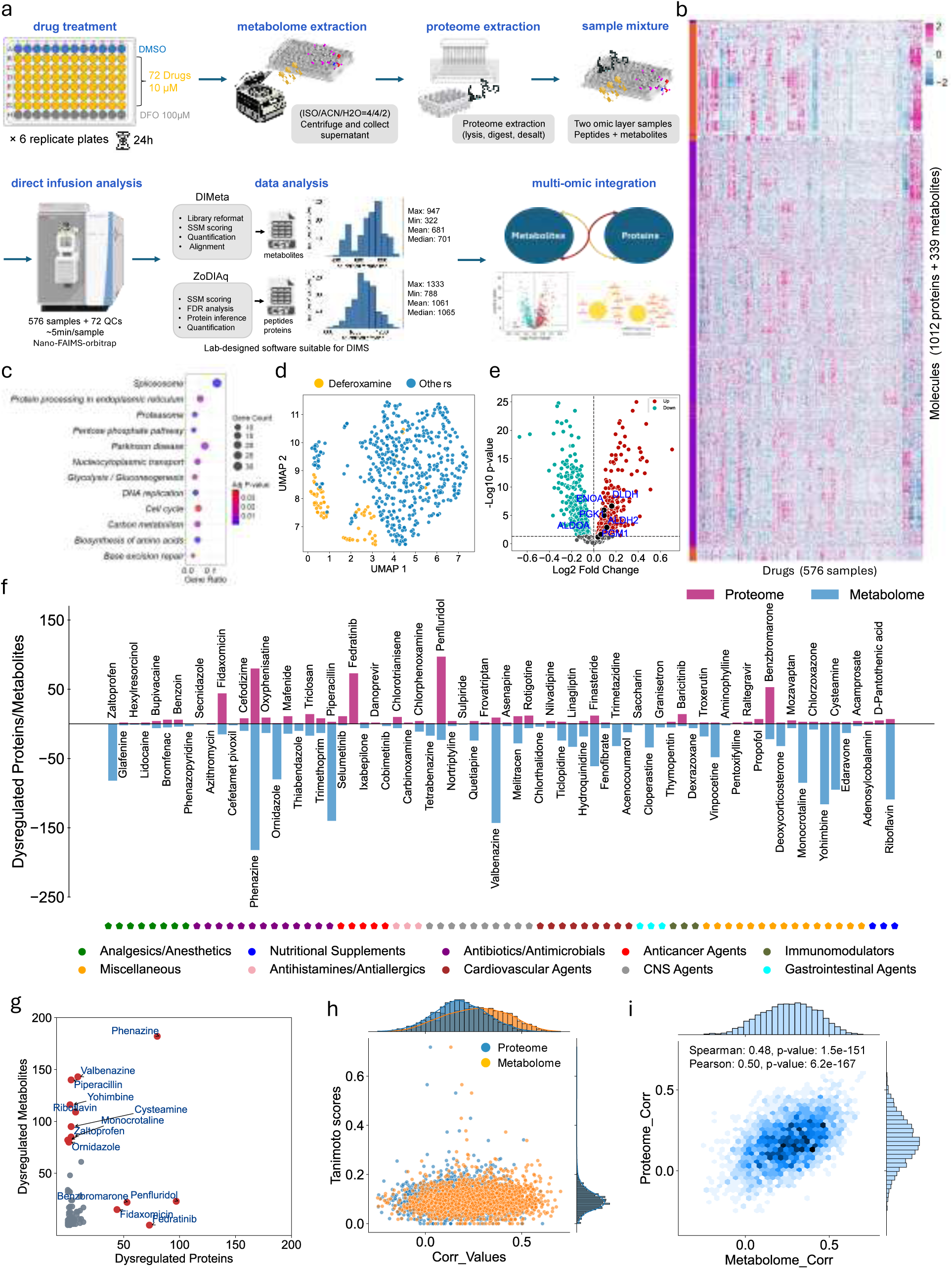
| SMAD enables integrative multi-omics strategies for large-scale drug screening. **a,** an overview of multi-omics drug screening strategy based on 96-well plate, from experimental design to data analysis. Multi-omics analysis in duplicate is performed after 24-h treatments. **b,** Heatmap of all quantified proteins and metabolites after dataset cleaning. **c,** KEGG enrichment result of proteome result. **d,** UMAP analysis of negative control drug DFO for proteome result. **e,** Volcano plot of dysregulated proteins after DFO treatment **f,** bar plot of dysregulated proteins/metabolites in each drug (Student T-test with Benjamini–Hochberg (BH) adjusted P values <0.05, n = 6 or 12). **g,** Activity of top drugs measured by the number of dysregulated proteins (x axis) and dysregulated metabolites (y axis). **h,** Relations of Pearson correlation and structure similarity (quantified by tanimoto scores) of drug pairs. **i,** metabolome correlation and proteome correlation of all drug pairs. (All data in this figure comes from 576 independent samples corresponding to unique wells in six 96 well plates. n=6 or 12 for drugs and control, respectively. Significance calculated by student T-test with Benjamini–Hochberg (BH) adjusted P values <0.05)

Notably, in our previous case studies, we observed that the metabolomic coverage in 96-well-based multi-omics analysis was relatively limited. Additionally, the identification workflow and speed of GNPS also constrain its applicability to large-scale sample analyses. To overcome this challenge, we substantially revamped our metabolomics identification pipeline, developing a novel software tool named DImeta, which is explicitly optimized for direct infusion metabolomics. DImeta facilitates metabolite identification through MS/MS spectral matching, combined with compensation voltage alignment, with quantification relying on the intensity of secondary fragment ions (**Fig. S14, Fig. S15a**). We designed a graphical user interface (GUI) for DImeta, incorporating customizable spectral library integration methods and identification parameters. The software also supports output of visualized spectral matching results for each identified metabolite, along with corresponding textual reports (**Fig. S15b-f**). This new tool significantly expedited both identification and quantification processes. In conjunction with Zodiaq for proteomic analysis, our enhanced workflow achieved robust identification, averaging 1,061 proteins and 681 metabolites per sample (**Fig. 5a**). In other words, our method can generate an integrated multi-omics dataset encompassing over one million feature measurements (576 samples × [1,061 proteins + 681 metabolites]) in less than three days.

To ensure data quality, we performed rigorous data preprocessing, including the removal of outliers, the removal of replicate features, the removal of missing values, and data normalization (**see methods**). Ultimately, 1,012 proteins and 339 metabolites were retained for downstream analysis (**Fig. 5b, Supplementary Table 4**). We first assessed the quality of this dataset. From a proteomic perspective, many key metabolic pathways, such as DNA replication, cell cycle regulation, and central carbon metabolism, were well represented (**Fig. 5c**). Dimensionality reduction analysis revealed that the negative control samples clustered tightly (**Fig. 5d**), indicating acceptable technical variation after data preprocessing. In addition, glycolysis-related proteins were significantly upregulated (**Fig. 5e**, **Fig. 12c,d**), consistent with the known effects of DFO treatment and our previous findings.^37^ On the metabolomics side, several major metabolic pathways were also detected, and control samples also clustered together, supporting the robustness of the dataset (**Fig. S12 e-g**). Quantified multi-omics features also exhibited a stable coefficient of variation (CV) for all drugs (**Fig. S12h**).

We then conducted differential analysis for each drug treatment, examining both proteomic and metabolomic changes relative to the controls (**Fig. 5f**). A key initial observation was that most compounds did not induce substantial multi-omics perturbations. Notably, metabolic changes were generally more pronounced than proteomic changes, which aligns with expectations, as most selected drugs were non-cytotoxic and small molecules tend to perturb metabolism more readily than protein expression (**Extended Data Fig. S12i,j**). Another important finding was that the number of significantly altered metabolites did not show a linear relationship with the number of altered proteins. For example, riboflavin extensively remodeled metabolites with little effect on proteins (**Fig 5f**). In other words, extensive perturbation at one omics level did not necessarily result in equally extensive changes at the other (**Fig. 5g**).

A third notable observation was that drugs belonging to the same pharmacological class did not necessarily induce similar levels of multi-omics alterations. This may be due to the broad nature of conventional drug classification systems, where compounds within the same category may act via distinct mechanisms or target different pathways. To explore this further, we incorporated a chemical similarity scoring system using the Tanimoto coefficient. The analysis revealed that most drugs exhibited low structural similarity, with Tanimoto scores below 0.2. The only pairs with relatively high similarity were VMAT2 inhibitors, Tetrabenazine and Valbenazine; however, even these did not show strong multi-omics correlation (**Fig. 5h**).

Although the number of significantly altered features in the two omics layers did not correlate after multiple testing correction, we observed a moderate linear relationship in the number of altered molecules when considering the uncorrected p-values (**Fig. S12k**). This prompted us to investigate further whether the direction and magnitude of changes across the proteome and metabolome were concordant. We calculated the Pearson correlation coefficients for proteomic and metabolomic profiles across all drug pairs, respectively. The results revealed a moderate correlation (R ≈ 0.5) between proteomic and metabolomic variation trends following drug treatments (**Fig. 5i**). This suggests that, although the number of significantly dysregulated features differs between omics layers, their directional changes and response magnitudes are aligned to some extent, indicating a potential intrinsic connection and coordinated regulation between the proteome and metabolome.

### Multi-omics perturbation drives the organization of drug communities

We further performed Pearson correlation analysis across all drug pairs based on their multi-omics features. Overall, we found relatively low correlations between most drug pairs, with the majority exhibiting correlation coefficients below 0.3. By applying a correlation threshold of 0.35, we constructed a drug–drug similarity network consisting of 469 edges. Among the 73 tested compounds (including DFO), 69 showed significant correlation with at least one other drug (**Fig. 6a**).

**Fig.6.**
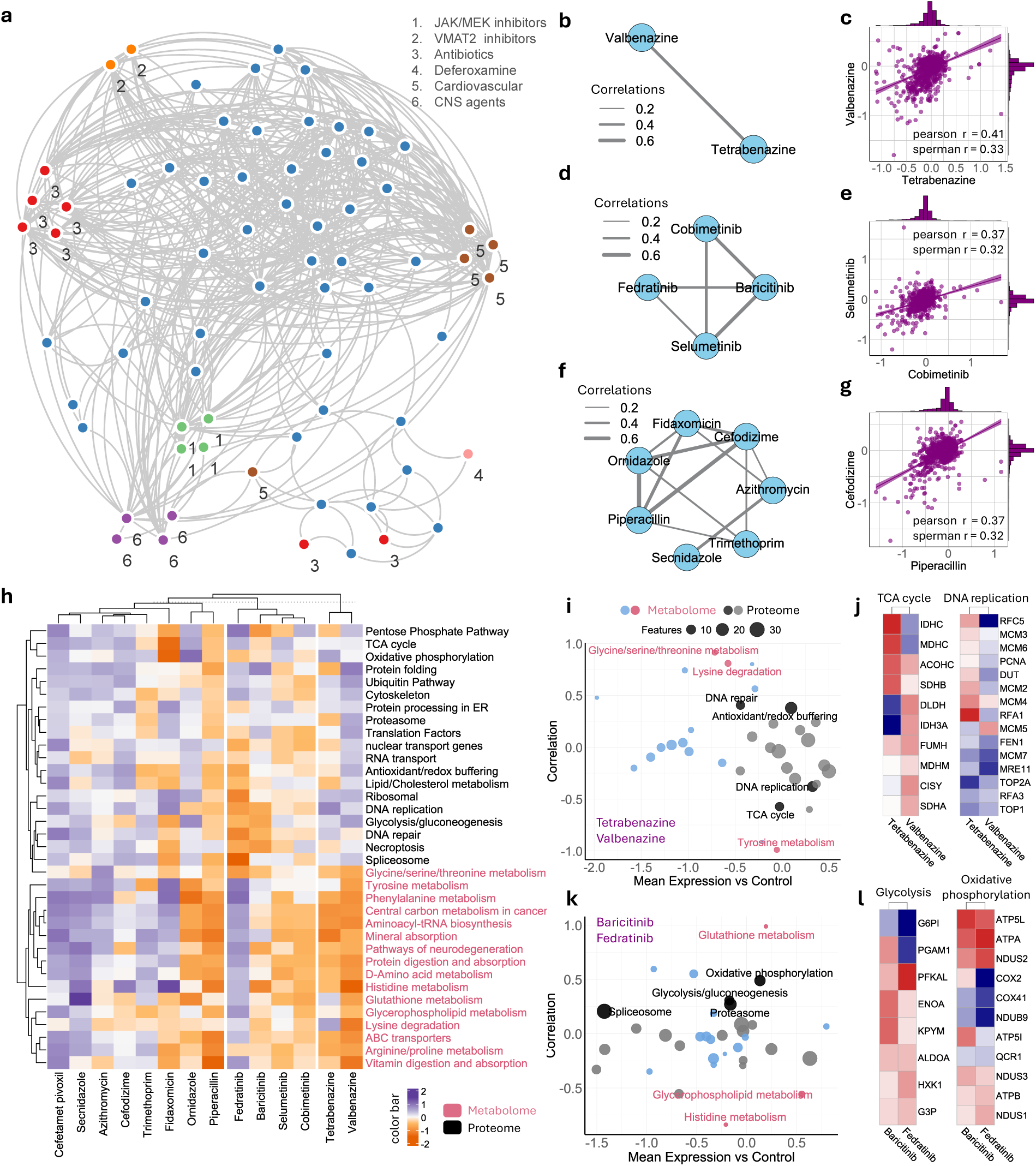
| Drug clustering driven by multi-omics response similarity. **a,** Community plot built from a compound–compound correlation matrix. Typical clusters were labeled with different color. Correlations are filtered to only include edges with r > 0.35. **b,c,** Subcommunity of VMAT2 inhibitors(b) and Pairwise correlation plot of VMAT2 inhibitors Valbenazine and Tetrabenazine(c). **d,e,** Subcommunity of JAK/MEK inhibitors(d) and Pairwise correlation plot of MEK inhibitors selumetinib and cobimetinib(e). **f,g,** Subcommunity of antibiotics(f) and Pairwise correlation plot of two typical antibiotics Cefodizime and Peperacillin(g). **h,** Clustered heatmap illustrates the changes in multi-omics pathways across different drug clusters, with metabolomics (red) and proteomics (black). **i,** Correlation and perturbation patterns of VMAT2 inhibitors (Valbenazine and Tetrabenazine) across proteomic and metabolomic pathways. **j,** changes in associated molecules within representative dysregulated pathway related to **i**. **k,** Correlation and perturbation patterns of JAK inhibitors (Baricitinib and Fedratinib) across proteomic and metabolomic pathways. **l,** changes in associated molecules within representative dysregulated pathway related to **k**.

Within this network, several distinct clustering patterns emerged based on multi-omics data, reflecting both structural and functional relationships among compounds. A notable example is the pairing of Tetrabenazine and Valbenazine, two structurally analogous inhibitors targeting the vesicular monoamine transporter 2 (VMAT2). Their substantial multi-omics similarity (**Fig. 6b,c**) is likely attributed to their shared molecular structures and analogous mechanisms of action, influencing comparable pathways and downstream molecular responses. Additionally, a distinct cluster comprising Janus kinase (JAK) and mitogen-activated protein kinase kinase (MEK) inhibitors was identified. Interestingly, despite significant structural diversity among compounds in this cluster, they converge on common signaling pathways involved in cellular proliferation, differentiation, and immune modulation. Consequently, these shared biological targets result in moderate omics-level correlations, reflecting a convergence of effects at the cellular and molecular level despite structural differences (**Fig. 6d,e**).

Furthermore, clusters based primarily on functional similarity were also observed. These include groups of antibiotics, central nervous system (CNS) agents, and cardiovascular drugs (**Fig. S16a,b**). Although grouped within the same therapeutic class and demonstrating some level of correlation (**Fig. 6f,g)**, these drugs frequently operate through varied molecular mechanisms, resulting in less pronounced multi-omics associations compared to the other two types of clusters. For instance, within the antibiotic cluster, compounds such as Secnidazole and Trimethoprim demonstrated notably weaker correlations with other antibiotics (**Fig. 6f, Fig. S16c**). A similar pattern was also observed in the CNS and cardiovascular drug clusters (**Fig. S16d-g**). This lower correlation could be attributed to their distinct modes of antimicrobial action, which target different metabolic or synthetic pathways within bacterial cells, thereby eliciting divergent molecular signatures detectable by omics profiling. These findings underscore the complexity and multidimensional nature of drug interactions, highlighting the need for integrated omics analyses to gain deeper mechanistic insights into the functionality and classification of drugs.

Given the complexity and scale of features in multi-omics data, direct correlation calculations may introduce noise and potential bias. To address this, we extracted a panel of 35 biological pathways, comprising 19 protein-based and 16 metabolite-based pathways (**Fig. 6h**), and conducted pathway-level comparisons across the three clustering types mentioned above. Interestingly, we found that drugs grouped into the same cluster at the global level also can demonstrate divergent or even opposing behaviors at the individual pathway level. For example, although Tetrabenazine and Valbenazine—two structurally similar VMAT2 inhibitors—displayed consistent trends across most metabolomic pathways, they showed striking differences in several proteomic pathways, including inverse patterns in TCA cycle and DNA replication (**Fig. 6i,j**).

Another example is the antibiotic Piperacillin, which exhibited similarity to Ornidazole in metabolomic pathways but differed significantly from other antibiotics (**Fig. 6h, Fig. S16i,m**). Likewise, Fidaxomicin, another antibiotic, demonstrated distinct profiles from the rest of the antibiotic cluster across both omics’ layers (**Fig. 6h**). Similarly, within the CNS drug cluster, Penfluridol exhibited substantial differences from other drugs in the group at the proteomic pathway level, whereas Quetiapine showed pronounced divergence from other cluster members at the metabolomic pathway level (**Fig. S16h,j**). A similar phenomenon was also observed within the cardiovascular drug cluster (**Fig. S16h,k**).

Notably, For the JAK and MEK inhibitors, we found that Baricitinib and Fedratinib (both JAK inhibitors) differed markedly in some metabolome-related pathways like histidine metabolism and Glycerophospholipid metabolism, yet were highly similar in key proteomic pathways such as oxidative phosphorylation and glycolysis (**Fig. 6k,l**). In contrast, Cobimetinib and Selumetinib, both MEK inhibitors, showed relatively better correlations in both proteomic and metabolomic dimensions (**Fig. 6h, Fig. S16l,n**). While pathway-level analyses may not fully capture the global multi-omics perturbation patterns—and may even miss critical information—they nonetheless underscore the complexity and diversity of cellular responses to drug treatments. Taken together, these results highlight the necessity of integrating multiple omics layers to understand drug mechanisms and their biological impact comprehensively.

### Exploration of intrinsic connections among multi-omics molecules

Given the extensive molecular interactions and co-perturbation phenomena observed both within individual omics layers and across multi-omics layers in our other case studies, we became deeply interested in further exploring the potential relationships among these multi-omics molecules. To this end, we first constructed a correlation network encompassing all multi-omics features, aiming to identify molecular entities that exhibit synergistic or antagonistic functions. Specifically, we calculated pairwise correlations among all molecules and selected those with correlation coefficients greater than 0.7. The resulting network consisted of 258 nodes and 1,249 edges.

Upon categorizing the molecules, we observed a tendency for features within the same omics layer to cluster together. However, a considerable number of molecules also demonstrated high correlations across different omics layers. Moreover, we found that multi-omics molecules involved in the same metabolic pathways or sharing similar biological functions were more likely to form clusters. For instance, molecules associated with glycolysis, cytoskeletal organization, and apoptosis—including both proteins and lipid/amino acid-related metabolites—tended to aggregate, which aligns well with biological expectations and further supports the robustness of our analytical approach (**Fig. 7a**).

**Fig.7.**
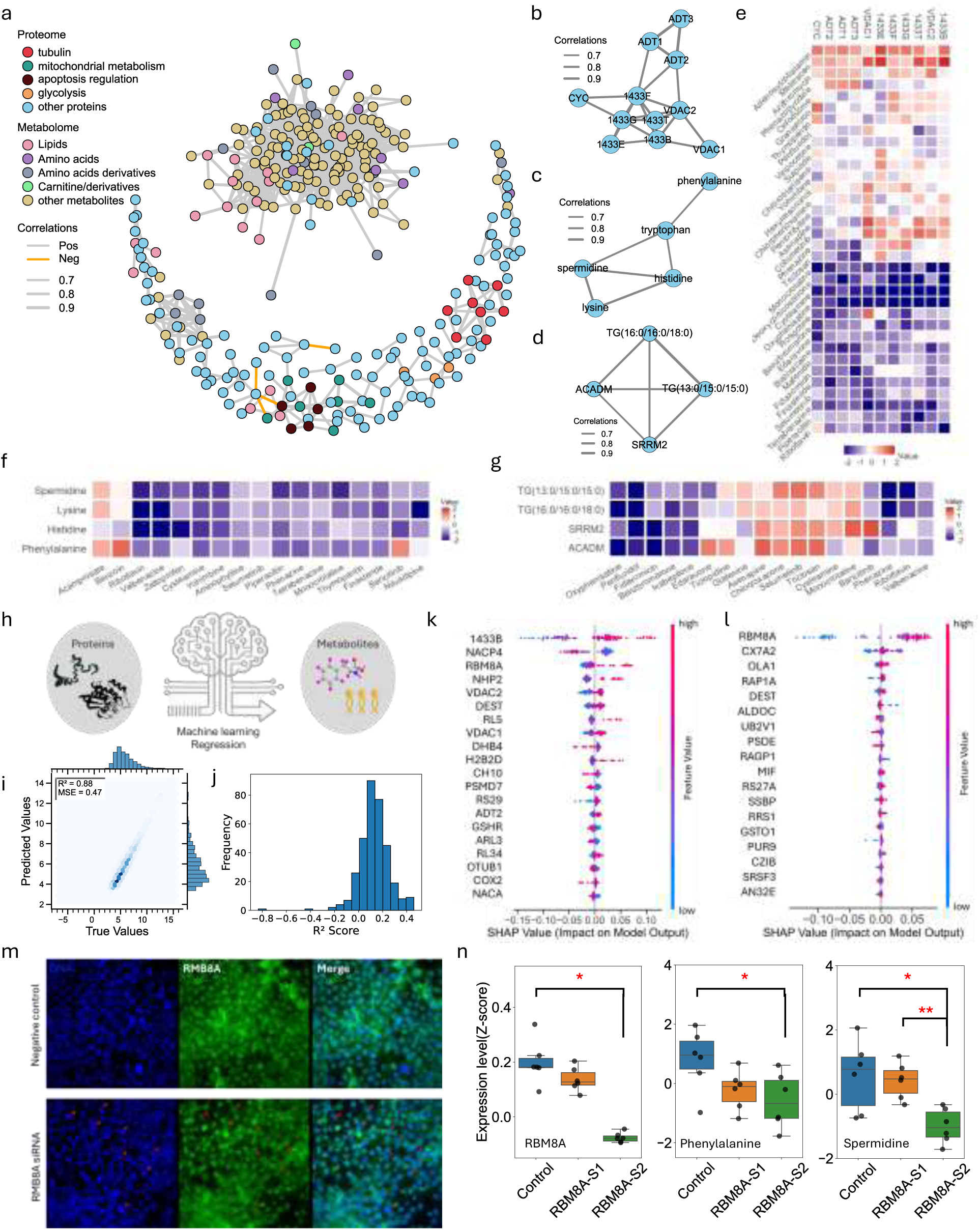
| Exploration of the intrinsic connections among multi-omics molecules. **a,** Community plot built from a molecule–molecule correlation matrix. Typical clusters were labeled with different color. Correlations are filtered to only include edges with r > 0.7. **b,** Subcommunity of typical proteome cluster. **c,** Subcommunity of typical metabolome cluster. **d,** Subcommunity of typical multi-omics interaction cluster. **e,** heatmap of selected protein features with higher correlation that significantly dysregulated their expression in compounds. **f,** heatmap of selected metabolite features with higher correlation that significantly dysregulated their expression in compounds. **g,** heatmap of selected multi-omics molecule features with higher correlation that significantly dysregulated their expression in compounds. **h,** Scheme showing the ML task of predicting metabolites from proteomic data measured in parallel by SMAD. **i,** Overall performance of all metabolites true versus ML predicted values. **j,** histogram of R squared values for all predicted metabolites. **k,l,** SHAP value of top contributed features of predicted molecule spermidine (k) and phenylalanine (l). **m,** Validation of RBM8A gene knockdown. **n,** (left) LC-MS results showing RBM8A level is significantly downregulated in the second knockdown group (p < 0.01), LC-MS metabolome analysis showing Phenylalanine level (middle) and Spermidine level (right) are significantly downregulated after RBM8A knockdown (p < 0.05). S1 and S2 represents two RBM8A siRNAs.

Further extraction of these highly correlated features revealed several intriguing sub-clusters. For example, significant correlations among the proteins ADT1, ADT2, ADT3, CYC, VDAC1, VDAC2, and the 14-3-3 protein isoforms 1433B, 1433E, 1433F, 1433G, and 1433T within cells, suggesting potential coordinated roles or shared pathways (**Fig. 7b**). The ADT proteins are involved in cellular energy metabolism and redox homeostasis, whereas VDAC family proteins function as mitochondrial membrane channels regulating the transport of metabolites and ions, thus directly influencing mitochondrial metabolism and apoptotic signaling. Cytochrome C (CYC) is a critical component of the mitochondrial respiratory chain, participating in both energy production and the induction of apoptosis. Additionally, the 14-3-3 family proteins are well-known modulators of various protein-protein interactions, cell-cycle progression, and apoptotic signal transduction pathways. The observed robust correlations among these proteins likely reflect a coordinated regulatory mechanism integrating metabolic activities, mitochondrial functions, and apoptotic signaling.

Moreover, our results also revealed strong correlations among the metabolites lysine, histidine, phenylalanine, tryptophan, and spermidine, suggesting an integrated metabolic network within the cell (**Fig. 7c**). These amino acids participate in diverse biological functions, including protein biosynthesis, metabolic signaling, and the regulation of oxidative stress. Spermidine, derived from amino acid metabolism, plays a crucial role in cellular proliferation and survival. The observed coordinated changes among these metabolites likely reflect shared regulatory mechanisms responding to cellular energy status, nutrient availability, or stress conditions. Although these metabolites don’t share a single linear pathway, they intersect metabolically at central hubs, such as the TCA and urea cycles.

More interestingly, the strong correlation among a set of multi-omics features was observed (**Fig. 7d**), including ACADM (medium-chain acyl-CoA dehydrogenase), triglycerides (TG(13:0/15:0/15:0), TG(16:0/16:0/18:0)), and SRRM2 (Serine/Arginine Repetitive Matrix 2). Functionally, ACADM is essential for fatty acid β-oxidation and may influence the metabolism of these triglycerides, while SRRM2 regulates pre-mRNA splicing. Their coordinated behavior suggests a potential link between lipid metabolism and post-transcriptional gene regulation. Fatty acid fluctuations could affect SRRM2 expression through energy or signaling pathways, whereas SRRM2-mediated splicing may impact the expression of lipid-related transcripts.

To further understand relationships between the metabolomic and proteomic data we used an unbiased machine learning approach similar to our previous work.^50^ We first test this idea with dataset from case study 2 to predict changes in proteins from changes in metabolites using an optimized extratrees model ^51^ (**Fig. 17a**). Overall, the prediction of most proteins was very effective with an R2 score of 0.7 between true and predicted values (**Fig. S17b**). Several proteins from diverse pathways were predicted with remarkable accuracy with R2 scores over 0.98 (**Fig. S17c-e**). Given the task of predicting proteins from metabolites, we were not surprised to see metabolic proteins predicted well. Still, we were surprised to see a ribosomal protein among the top well-predicted proteins (**Fig. S17e**). In fact, the distribution of all proteins predicted from metabolites indicates that most proteins were predicted with rho over 0.8. Most ribosome proteins in our dataset were well predicted (**Fig. S17f**). To determine which proteins were statistically significantly well predicted by machine learning from metabolites measured from the same sample by SMAD, we used a strict Bonferroni correction of the p-values from Spearman correlation between the true and predicted protein values. This left us with 54 statistically well-predicted proteins. Term enrichment analysis of the most well-predicted proteins revealed that diverse pathways beyond metabolic proteins are accurately predicted from the metabolomic data, indicating strong relationships between the metabolome and multiple cellular pathways, including protein synthesis and degradation. This makes sense because most cellular energy is used for protein regulation.^52^

Encouraged by these results, we further applied the Random Forest Regressor model to predict metabolites from proteins (**Fig. 7h**). Overall, the prediction of most metabolites was very effective with an R square score of 0.88 between true and predicted values (**Fig. 7i**). The per-metabolite overall R2 distribution showed that some metabolites were predicted better than others (**Fig. 7j**). Spermidine and phenylalanine were among the best predicted metabolites (R^2^ score 0.41 and 0.42), so we used SHAP to determine which proteins were responsible for their accurate prediction (**Fig. 7k, 7l**). This revealed that no expected proteins known to regulate these metabolites were identified as regulators; however, RBM8A was found to be important for both metabolites. RBM8A is a core component of the exon junction complex, which is crucial for the post-transcriptional regulation of mRNA. It is known to help degrade mRNAs that contain premature stop codons and to influence mRNA localization. We used siRNA to knockdown GAPDH (control, **Fig. S18**) or RBM8A, one of the two probes employed (S1 and S2), resulted in a reduction of RBM8A (S2), as determined by both immunofluorescence and proteomics (**Fig. 7m, n**). As predicted by the SHAP, the loss of RBM8A in the S2 condition was associated with a drop in both phenylalanine and spermidine (**Fig. 7n**). This validates that even lower depth multi-omics from SMAD can predict causal relationships across omics layers.

## Discussion

Here, we present a high-throughput direct infusion mass spectrometry strategy, enabled by gas-phase ion mobility separation, that allows for the single-injection analysis of peptides, polar metabolites, and lipids in a single run. By integrating this method with our custom-developed software tools, we successfully identified over 1,300 proteins and 600 metabolites within approximately 4.3 minutes of data collection per sample. This workflow eliminates the need for liquid chromatography, significantly simplifying instrumental requirements, increasing sample throughput, and reducing analysis time. The successful implementation of this method is grounded in two key advancements: (1) gas-phase separation of peptides, lipids, and polar metabolites using both ion mobility and quadrupole selection, which effectively reduces ion complexity before detection and improves analyte coverage across omics layers; (2) tailored computational tools, such as ZoDIAq and DImeta, that support robust spectral matching and multi-omics integration.

We demonstrated the applicability and versatility of this method in three contexts. In macrophage polarization experiments, our approach captured treatment-induced multi-omics changes across several levels from global omics reprogramming and coordinated pathway regulation to individual protein-level alterations not previously reported. In a large-scale drug screening study, we observed both convergent and divergent omics responses across structurally and functionally related compounds, suggesting that global multi-omics profiles can reveal both shared mechanisms of action and drug-specific effects.^53, 54^ Furthermore, by incorporating machine learning–based integrative analysis, we identified cross-omics interaction patterns—e.g., proteo-metabolomic correlations—which were validated using biological assays and conventional LC-MS-based platforms.

Despite its advantages, the current SMAD approach still falls short of traditional LC-MS methods in terms of depth and dynamic range of identification. Moreover, the limited resolving power of FAIMS leads to residual overlap between the two omics layers, which in turn can compromise identification efficiency. Furthermore, our workflow relies on label-free quantitation—an approach widely used in conventional LC–MS—but performing single-injection multi-omics by direct infusion may exacerbate matrix effects and thus further undermine the accuracy of label-free quantitation. However, despite these limits, we believe this method still falls short of its theoretical ceiling. To push the boundaries of performance, advancements are needed across both hardware and software dimensions: (1) Ion Mobility: While FAIMS offers partial ion separation, integrating more powerful techniques such as trapped ion mobility spectrometry (TIMS) or structures for lossless ion manipulation (SLIM) could further enhance gas-phase resolution and reduce chimeric spectra, especially across distinct omics layers. (2) Mass analyzer: The Orbitrap’s detection speed (∼100–150ms per scan) presents a throughput bottleneck. Employing faster platforms such as time-of-flight (TOF) or the newly emerging Astral analyzers^16, 55^ could substantially shorten analysis time, potentially enabling sub-minute total acquisition. (3) Ion Source Optimization: Improved ionization efficiency—through refined emitter geometries and ultra-low flow regimes—can enhance signal intensity while reducing sample consumption, both of which are critical for clinical and precious sample applications. On the computational side, our current spectral matching relies on cosine similarity. Future development of deep learning–based spectral deconvolution and annotation tools may yield higher identification sensitivity, better handling of mixed spectra, and improved cross-omics feature alignment.^26, 56, 57^

Looking forward, we recognize the need to reevaluate how this technology is positioned. Rather than universally striving for maximum omics depth, more impactful applications may lie in focused, clinically relevant use cases, such as rapid profiling of well-established biomarkers or pathway-specific panels in disease diagnostics. In these contexts, a platform that is simple, robust, fast, flexible, and cost-effective may provide greater clinical utility than one designed for maximal omics coverage. For instance, clinical assays often require the detection of just dozens of disease-associated metabolites or a few hundred proteins, with high reproducibility and minimal sample preparation.^58^ Similarly, SMAD may offer unique advantages in structural or interaction proteomics,^59^ enabling rapid gas-phase characterization of selected targets without the confounding influence of LC-based retention effects.

In summary, we demonstrate for the first time the feasibility of LC-free, single-injection multi-omics profiling using DI-MS. Coupled with integrated data processing and cross-omics analysis, this method offers a powerful platform for high-throughput cellular phenotyping, mechanistic studies, and pharmacological screening. We anticipate that ongoing advances in sample processing, instrument design, and computational analytics will further elevate the utility of SMAD in both research and clinical domains.

## Methods

### Materials

Angiotensin I (Sigma, A9650-1MG), QCAL Peptide Mix (Sigma, MSQC2) and Hela digest standard (Thermo Fisher Scientific, Catalog number: 88328) were dissolved into different concentrations with 50% acetonitrile (ACN) in 0.2% formic acid (FA). Lipid standards (product No. 330707) was purchased form Avanti. Drugs including deferoxamine mesylate salt (Product No. D9533), mTOR inhibitor torin2 (Product No. SML1224), integrated stress response inhibitor ISRIB (Product No. SML0843), proteasome inhibitor MG-132 (Product No. 474790) and SCD1 inhibitor A939572 (Product No. SML2356) were purchased from Sigma-Aldrich.

### Mass spectrometry and data acquisition

**SMAD** analysis of case study 1 and 2 was performed on an Orbitrap Lumos (Thermo Fisher Scientific) mass spectrometer coupled with the FAIMS Pro Interface. SMAD analysis of case study 3 was performed on an Orbitrap Ascend (Thermo Fisher Scientific) mass spectrometer coupled with the FAIMS Pro Interface. Different compensation voltages were applied for metabolome (−5V to −40V in steps of 5V) and proteome (−30V to −80V in steps of 10V) analysis. A nano-ESI source (“Nanospray Flex”) and LOTUS nESI emitters from Fossiliontech were used for ionization. The ultimate 3000 HPLC system (Thermo Fisher Scientific) was used to control automated sample loading, flow rate, and mobile phase composition. Flow rate was maintained at 1.4ul/min at the first 0.5 min for transferring samples to nano emitter and then maintained a 0.3ul/min flowrate to the end of the acquisition. Mobile phase composition is ACN/H2O (70:30) in 0.1% formic acid (FA) for the whole acquisition process. Data acquisition was conducted at positive mode with 2200V. AGC was set at 100% and ion injection time was set at auto. For proteome acquisition, targeted MS2 mode was used for each compensation voltage from −30 V to −80 V in a step of 10V. For metabolome acquisition, tSIM mode with a quadrupole window of 50da was used to scan full *m/z* range from 100 to 1100 for each compensation voltage from −5V to −40 V in a step of 5V. For case studies 1 and 2, three QC samples were applied every ten runs; for case study 3, twelve QC samples were applied between each plate for batch-effect correction.

**LC-MS/MS** proteomic analysis was conducted on an Orbitrap Ascend Tribrid Mass Spectrometer using data-independent acquisition (DIA) with the following settings: positive ionization (2500 V), 100 ms ion injection time, 12 Da DIA window, MS2 range 400–1000 m/z, AGC target 100%, and HCD energy at 30%. Peptides were first trapped on an EXP®2 Stem Trap and separated on a 200 cm µPAC™ column (PharmaFluidics), connected via 20 µm i.d. Viper™ capillaries (Thermo Fisher) and maintained at 55 °C. The reversed-phase gradient (A: 0.1% formic acid in water; B: 0.1% formic acid in acetonitrile) was as follows: 8% B to 25% at 70 min, to 37% at 95 min, ramped to 98% at 96 min (held for 9 min), then returned to 8% at 105 min and held for 5 min (total 110 min). For library construction, LC-MS/MS proteomic analysis was performed on an Orbitrap Lumos Mass Spectrometer using data-dependent acquisition (DDA) with the following settings: FAIMS compensation voltages ranging from –30 V to –80 V in 5 V increments (one compensation voltage per run, totaling 11 runs), positive ionization mode (2500 V), ion injection time of 100 ms, MS2 scan range of 300–1000 *m/z*, AGC target set to 100%, and HCD collision energy at 30%. The LC column and mobile phase settings were the same as those used in the DIA method described above.

**LC-MS/MS** metabolomic analysis was conducted on an Orbitrap Exploris 480 Mass Spectrometer using targeted MS/MS (tMS2) with the following settings: positive ionization mode (2600 V), ion injection time of 100 ms, MS2 scan range of 70–1000 *m/z*, AGC target set to 100%, and stepped HCD collision energy (15, 30, 45%). Metabolites were trapped and separated on a 50mm UPLC HSS PFP column (Acquity, 1.8 µm) maintained at 35 °C. The reversed-phase gradient (A: 0.1% formic acid in water; B: 0.1% formic acid in acetonitrile) was as follows: 2% B to 50% at 15 min, ramped to 98% at 30 min (held for 5 min, total 35 min).

### Quantitative Evaluation of SMAD

Three lipid standards (d18:1-18:1(d9) SM, 29.6 μg/ml; 15:0-18:1(d7) PC, 150.6 μg/ml; 18:1(d7) Lyso PC, 23.8 μg/ml) and QCAL proteins (0.25 μg/μl) was mixed and diluted every four times. The mixed standard sample was directly analyzed by targeting their accurate *m/z* with SMAD. Lipid standards and MS-QCAL peptides were quantified with python by manually extracting MS1 intensity and representative y-ion fragments intensity, respectively. For real samples, original metabolome samples were produced by adding 500ul metabolite extraction solvent (ISO/ACN/H2O, 4:4:2) to 2 million 293T cells and then the supernatant was collected. Proteome samples was derived from 293T cells with same proteome preparation protocol used for two case studies. Then the multi-omics sample was produced by mixture of metabolome and proteome samples at 1:1 volume ratio. The sample was diluted every four times to produce concentration gradients and analyzed by SMAD.

### Cell culture and sample preparation for macrophages

BMDMs were derived from bone marrow extracted from the femurs of euthanized mice (11-week-old male C57BL/6J) and plated at 3 × 10^6^ cells per 10-cm dish in 10 ml of macrophage growth medium (complete RPMI containing 25% M-CSF containing L929-conditioned medium). Cells were cultured for 7 days to differentiate and were supplemented with 5 ml of macrophage growth medium on day 5. On day 7, 7×10^6^ BMDMs were counted and replated on 10 cm dishes in macrophage growth medium overnight prior to experiments. On the day of experiments, BMDMs were treated with 100ng/ml LPS (Invivogen, LPS-EK Ultrapure) and 10ng/ml recombinant murine IL-4 (Peprotech) for 24 hours. Then the cells were washed twice with cold PBS and harvested from the plate by scraping. The cells were pelleted into 1.5 ml centrifuge tubes and snap frozen. Then the metabolites and lipids were extracted from samples with 500μl isopropanol/acetonitrile/water 4:4:2 for 20 min, following by a hard spin of 10 minutes in 10000 rcf and all metabolome supernatant was removed to new centrifuge tubes and stored in −80℃. The precipitated proteome pellet was dry out and then lysed by addition of 8 M urea with 50 mM TEAB buffer at pH 8.5. The tubes were vortexed and sonicated until homogenous with lysis buffer. Then TCEP and chloroacetamide were each added to 10 mM final concentration to reduce protein disulfide bonds and alkylate the free cysteines in the dark for 30min. Then lysis buffer was diluted to 2 M urea using 50 mM TEAB, and catalytic hydrolysis of proteins was initiated by trypsin (Promega) at a weight ratio of 1:50 protease:substrate. Proteome proteolysis was incubated in a 37℃ incubator for six hours. Peptides were desalted using Strata reversed-phase cartridges from Phenomonex, and then dried completely in Speed-Vac. Peptides were resuspended with ACN/Water/FA (50%/49.9%/0.1%, volume ratio) and mixed with metabolome samples for SMAD analysis. The final loading concentration of proteome is around 0.3ug/ul.

### Cell culture, sample preparation for drug screening, and gene knockdown

HEK293T cells were cultured in a 96-well plate to 70% confluency and then treated with different drugs. For case study 2, the treated concentrations of drugs were deferoxamine (10μM), Torin2 (1μM), ISRIB (1μM), MG132 (1μM) and A939572 (1μM). For case study3, the treated concentrations of all drugs are 10μM. Drugs were dispensed to 96 well plates with Echo 650 Series Liquid Handlers. After 24 hours incubation, cell culture media was removed and washed with PBS for twice. Then 80ul metabolite extraction solvent (IPA/ACN/H2O, 4:4:2) was added to each well of the plate and vortexed for ten minutes. After vertexing, the plate was centrifuged in 2,000 rcf and all metabolome supernatant was removed to a new 96-well plate and stored in −80℃. The precipitated proteome pellet in original 96-well plate was dry out first and then lysed by addition of 8 M urea with 50 mM TEAB buffer at pH 8.5. The plate was vortexed at 900 rpm around 5 minutes until homogenous with lysis buffer and sonicated 5 min in a Covaris sonicator (R230 focused-ultrasonicator) maintained at 4°C. After sonication, TCEP and chloroacetamide were added to a 10 mM final concentration to reduce protein disulfide bonds and alkylate the free cysteines in the dark for 30 min. Then, lysis buffer was diluted to 2 M urea using 50 mM TEAB, and catalytic hydrolysis of proteins was initiated by trypsin (Promega) at a weight ratio of 1:50 protease:substrate. Then Proteome proteolysis was incubated in 37℃ incubator for six hours. Peptides were desalted using 96-well µElution Plate from Waters (Oasis HLB 96-well µElution Plate, 2 mg Sorbent per Well, 30 µm), and then dried completely in Speed-Vac. Peptides were resuspended in 10 ul ACN/Water/FA (50%/49.9%/0.1%, volume ratio) and mixed with 10ul metabolome samples for SMAD analysis. The final loading concentration of proteome is around 0.15ug/ul. To prepare the 293T proteome for experimental parameter analysis, we followed a consistent protocol for cell lysis and digestion except that cells were cultured in a 12cm plate and subsequently desalted using Strata reversed-phase cartridges from Phenomonex. HEK293T cells were seeded in 24-well plates to reach 60% confluency at the time of transfection. Gene knockdown of GAPDH and RMB8A was performed using Silencer Select siRNAs (Thermo Fisher Scientific; Negative Control siRNA, Cat#4390843; GAPDH siRNA, ID#AM4624; RBM8A siRNA, ID# 137865 and 45102) at a final concentration of 10 nM per siRNA. Lipofectamine RNAiMAX transfection reagent was used to deliver siRNAs according to the manufacturer’s protocol. Knockdown efficiency was assessed 48 h post-transfection by either immunoblotting, immunofluorescence, or mass spectrometry.

### Polarization and generation of senescent BMDM culture

BMDMs used in experiments were derived from bone marrow extracted from the femurs of euthanized mice (6-12 weeks old male C57BL/6J) by mortar and pestle. Femurs were placed in the mortar and were washed with 70% ethanol to sterilize followed by two washes with complete RPMI (cRPMI; standard RMPI (Corning) supplemented with 10% fetal calf serum, penicillin-streptomycin solution (Corning), 1mM sodium pyruvate solution (Corning), 2 mM L-Glutamine solution (Corning), 10nM HEPES buffer (Corning), and 50μM 2-mercaptoethanol). After washing, 10 mls of cRPMI were added to the mortar and the femur bones were gently crushed. The resulting media was collected and filtered through a 70μm filter and placed in a conical tube. The filtered supernatant was centrifuged at 1200 RPM (150 RCF) for 5 minutes. Cells were resuspended, counted and plated at a density of 3E6 cells/10 cm dish in 10 ml of macrophage growth media (cRPMI containing 25% M-CSF containing L929 conditioned media (made in house)). Cells were left to grow for 7 days to differentiate and were supplemented with 5 ml of macrophage growth media on day 5. On day 7, BMDMs (yielding 10-12×10^6^ cells/10cm dish) were lifted off the plate using cold PBS containing 5mM EDTA. BMDMs were counted and replated in macrophage growth media overnight prior to experiments. On day of experiments, macrophage growth media was replaced with cRPMI 6 hours prior to stimulation to remove M-CSF. M2 polarization was performed by stimulating macrophages with 10 ng/ml recombinant mouse IL-4 (Peprotech). For M1 polarization macrophages were stimulated with 100 ng/ml LPS (LPS EK-Ultrapure, Invivogen). On day 7, BMDMs were irradiated (10 gy) for 24 hours at 250 nM in 60% cRPMI: 40 MGM. DNA damaged cells were left in culture for 10 days and media components were replaced every 2-3 days.

### Proteome and metabolome library generation

The building of proteome library used for ZoDIAq generally including three steps: 1) performing LC-MS/MS analysis of Macrophage and 293T samples with DDA for eleven compensation voltages from −30V to −80V in a step of 5V. 2) Using Fragpipe to produce pepxml files of peptides and proteins with appropriate fasta database and add decoys (50%). 3) Building library for ZoDIAq with SpectraST. The latest version of ZoDIAq also supports proteome library generated from Fragpipe.

The building of metabolome library used for DImeta including two steps:

1. The Orbitrap-based standard metabolite spectral library was downloaded from GNPS website (https://external.gnps2.org/gnpslibrary).
2. The spectral data were processed and integrated into a final consensus library using the “Library handling” module of the DImeta software. The resulting metabolite spectral library used in this study has been uploaded to the public database.

### Identification and quantification of proteome

Peptides and proteins were identified with ZoDIAq (https://github.com/xomicsdatascience/zoDIAq). ZoDIAq is a python software package designed to enhance usability and sensitivity of the projected spectrum−spectrum match scoring concept. The process was described as the following steps: Firstly, the original Thermo .RAW files were converted to mzXML files using msconvert with default settings. Then the output mzXML files were input to ZoDIAq GUI and the appropriate spectral library for that sample was selected with the default settings keeping the fragment mass tolerance at 20ppm and the “Label free quantification” should be selected. ZoDIAq produces three output files for each input mzXML file that report spectra, peptides, and proteins filtered to <1% FDR. In each case, ZoDIAq sorts peptide identifications by match count and cosine (MaCC) score, calculates the FDR for each identification using a modification of the target-decoy approach where FDR at score S = number of decoys/number of targets, and removes SSMs below a 0.01 FDR threshold. The peptide FDR calculations only use the highest-scoring instance among all SSMs for each peptide. ZoDIAq uses the IDPicker algorithm to identify protein groups from the list of discovered peptides and adds them as an additional column in the output. A detailed description about data processing, FDR calculation and protein inference was listed in our previous paper^26^. Peptides and proteins were quantified using ZoDIAq to extract the sum of all detected fragment ion intensities of common peptides in all input files. It is worth noting that, owing to FAIMS’s limited resolving power, a very small number of MS/MS spectra acquired at intermediate compensation voltages may contain fragment ions from metabolites; however, these occurrences are exceedingly rare—primarily below *m/z* 300—and do not compromise ZoDIAq’s identifications of proteome. DIANN 1.9.2 was applied for LC-MS proteome results analysis.

### Identification and quantification of metabolome

For Case Study 3, the steps for metabolite identification and quantification are as follows: Thermo .RAW files were converted to mzML files using msconvert. The produced mzML files were analyzed with self-developed DImeta software (https://github.com/xomicsdatascience/DImeta). Metabolome library was standard metabolite MS2 spectral downloaded from GNPS (https://external.gnps2.org/gnpslibrary). The downloaded libraries were integrated and standardized using the database operation module of DImeta. DImeta supports MS/MS spectral matching for metabolite identification, inter-sample alignment, and quantification. The output is provided in .csv format, with entries including precursorMZ, compensation voltage, cosine_score, ion_count, scan_number, compound_name, compoundMZ, adduct type, compound_formula, matched_peaks and macc_score. For other studies, Thermo .RAW files were converted to mzML files using msconvert. All peaks are picked according to the sequence of Q slices. The produced mzML files were analyzed with MZmine3 software for mass detection, feature detection, alignment, gap filling and feature filter. The output files contain *m/z* and quantification by FAIMS peak area. For metabolite annotation, we applied a direct infusion based DDA tandem mass spectrometry analysis with the same sample and same compensation voltages applied in SMAD analysis. Then the MS2 spectrum of metabolites was compared with library and analyzed through GNPS website.^22^ Finally, the identified metabolites and their corresponding accurate molecular masses were compiled into a database, which was then used for subsequent identification of the same type of samples.

### Consensus clustering and dimensionality reduction

In two applications of macrophage polarization and drug screening, the unsupervised k-means consensus clustering of all treatments was performed with the python packages “sklearn”. The significantly dysregulated molecules that were discovered among different treatments were used for clustering. The number of groups for clustering was determined by “Elbow Method”. PCA and UMAP analysis was performed with the python packages “sklearn” and “UMAP”, respectively.

### Pathway enrichment analysis

The UniProt IDs from ZoDIAq outputs were converted to gene IDs. The KEGG_2022_Human gene set library was applied and pathway enrichment analysis was done in Cytoscape ^60^ with the plugin clueGO.^61^ GO Term/Pathway network connectivity (Kappa score) was set at 0.5. Additional settings included GO Term grouping and two-sided hypergeometric tests, and leading group term ranking based on highest significance.

### Data Analysis

In case studies 1 and 2, Data preparation, analysis, and visualization were performed in Python version 3.9.7. multi-omic features more than one-third missing values across all treatments were removed. KNN-imputer was applied for missing value imputation after log2 transformation. One-way ANOVA was applied for group analysis and select the significant dysregulated molecules (Benjamini–Hochberg (BH)-adjusted P values <0.05) among control and treatments. two-sided Wilcoxon rank-sum test was used to compare each treatment with control and significant features were defined as a p-value less than 0.05 after Benjamini–Hochberg (BH)-adjustment. K-means clustering was applied for dysregulation pattern clustering after z-score normalized.

In case study 3, data preparation, analysis, and visualization were performed in Python version 3.9.7. Multi-omic features with more than half missing values across all drugs were removed. A KNN-imputer was applied for missing value imputation after total counts were normalized. ComBat was applied for batch effect correction. A Student t-test was used for dysregulation analysis between each treatment and control, and significant features were defined as those with a p-value less than 0.05 after Benjamini–Hochberg (BH) adjustment. Pearson and Spearman correlation analysis were applied for drugs and multi-omics features correlation analysis.

### Model Training and Evaluation

For the small test dataset of one 96 well plate, machine learning was performed using scikit-learn in python. Data was split into training and test sets where one sample from each of the 7 conditions was stratified into the test set. An extratrees model (n_estimators = 100, max_features = “auto”, max_depth = None, min_samples_split = 2, min_samples_leaf = 1, max_features = 1, bootstrap = False, max_samples = None) was optimized on the training data using 5-fold cross validation. A single model to predict each protein was trained using the best parameters from 5-fold cross validation and then the model performance was evaluated by predicting the quantity of that proteins in the test set by computing the mean squared error (MSE), R² scores and Spearman’s rank correlation.

For the large drug screening dataset, data were split into training and testing sets using GroupShuffleSplit, ensuring that all instances associated with a given drug remained in the same partition. A Random Forest Regressor model was trained using the protein expression dataset as input features and metabolite levels as the target variables. The model was optimized using default hyperparameters, and predictions were made on the held-out test set. Model performance was assessed using the coefficient of determination (R² score), which quantifies the proportion of variance explained by the model.

### Feature Importance Analysis Using SHAP

To interpret the impact of individual protein features on metabolite predictions, feature importance was assessed using SHapley Additive exPlanations (SHAP). SHAP values were computed using a Kernel Explainer, which estimates feature contributions based on their impact on model predictions. To illustrate feature importance for specific cases, the phenylalanine and spermidine models were selected for in-depth SHAP analysis. SHAP values were computed for each model, and beeswarm plots were generated to identify key protein features influencing phenylalanine or spermidine levels.

## Data Availability

The data that support the findings of this study are openly available from massive.ucsd.edu at https://doi.org/doi:10.25345/C5BV7B77J, reference number MSV00009841.

## Code Availability

Python version 3.9.7 was used and all code for multi-omics analysis of two case studies, quantification evaluation, and data visualization are provided open source via xomicsdatascience github https://github.com/xomicsdatascience/SMAD-project.

## Declaration of generative AI and AI-assisted technologies in the writing process

During the preparation of this work the authors used GPT models to refine the wording. After using this tool/service, the authors reviewed and edited the content as needed and take full responsibility for the content of the publication.

## Acknowledgements

We thank Dasom Hwang for their help with graphic design. We thank the NIH for funding (R35GM142502, R35GM156893, and R21AG074234).

## Author contributions

Y.J., and J.G.M. conceived and designed the study; I.S.P and A.J.C prepared macrophages samples. Y.J. prepared all other samples and acquired multi-omics data; Y.J performed the performance analysis, quantification analysis and multi-omics analysis of two case studies. A.M and J.G.M performed machine learning study. J.M.E performed validation experiments. Y.J. and J.G.M. prepared all the figures and wrote the manuscript.

