## Supplementary figures for "Single-Injection Multi-Omics Analysis by Direct Infusion Mass Spectrometry"

### for

### Supplemental Methods

**Proteome library build:**

Procedures:

1. LC-MS/MS analysis with data dependent acquisition (DDA)

• Perform LC-MS/MS analysis of experimental samples with DDA for same compensation voltages you plan to use for subsequent DISPA experiments. Here we used 11CVs from -30V to -80V in a step of 5V for T293 proteome analysis.

• Tip: Extra compensation voltages or repeated injections can be collected if you want a better coverage in your library.

2. Get peptide identifications in pep.xml files from FragPipe

• Download Fragpipe from https://fragpipe.nesvilab.org/ and install.

• Using Fragpipe to produce pepxml files of peptides and proteins. Instructions of FragPipe can also be found on the above website.

• Select appropriate initial reference fasta database for analysis. The initial reference fasta database we used here for 293T library build is 2022-08-02-decoys-reviewed-contam-UP000005640.fas, File contains 40840 entries (20420 decoys: 50.0%).

3. Produce spectra library files with SpectraST.

• Here we use SpectraST, a spectral library building and searching tool, to build library for CsoDIAq software. The SpectraST library creation commands we used are provided as a file in supplementary .txt file

• Details for commands of SpectraST can be found in website as below:

http://tools.proteomecenter.org/wiki/index.php?title=Software:SpectraST#Creating_Libraries_from_Sequence_Search_Results

**Metabolome library build:**

Procedures:

1. Download standard metabolite MS2 spectral library build on Orbitrap from GNPS (https://external.gnps2.org/gnpslibrary)

2. Using DImeta module to integrate and format libraries to a final output library for identification

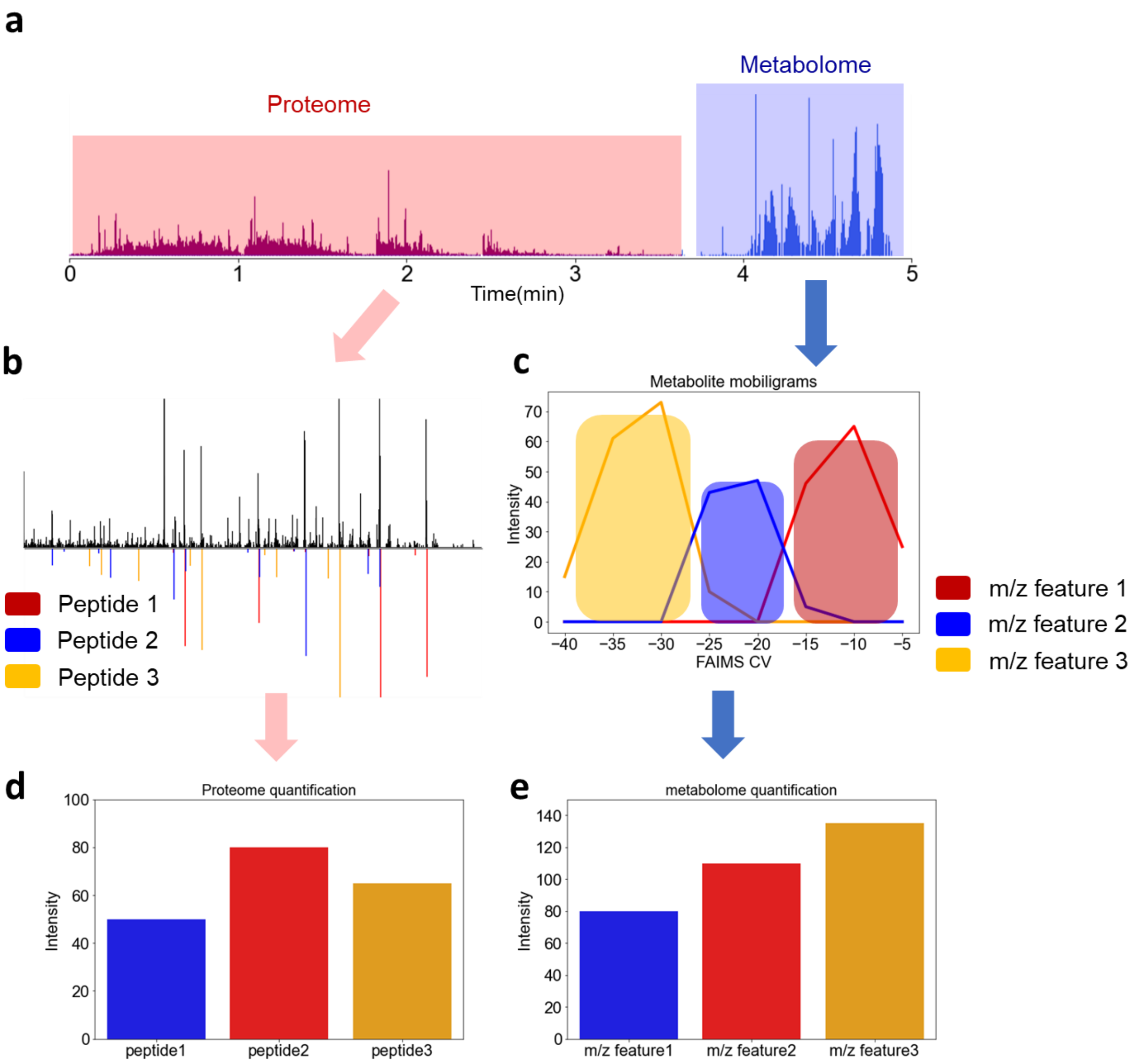


**Figure S1. Quantitative and identification schematic diagram of SMAD. a,** Typical TIC figure of SMAD. **b,** identification of peptides from tandem mass spectra by spectra-to-spectra match. **c,** extracting intensities with same m/z across all FAIMS compensation voltages and calculate ion mobiligrams (XIMs) for quantification. **d,** Quantification of proteome by summing up all fragment intensities of a specific peptide. **e,** Quantification of metabolome by extracted ion mobiligrams (XIMs) of a specific *m/z* feature.

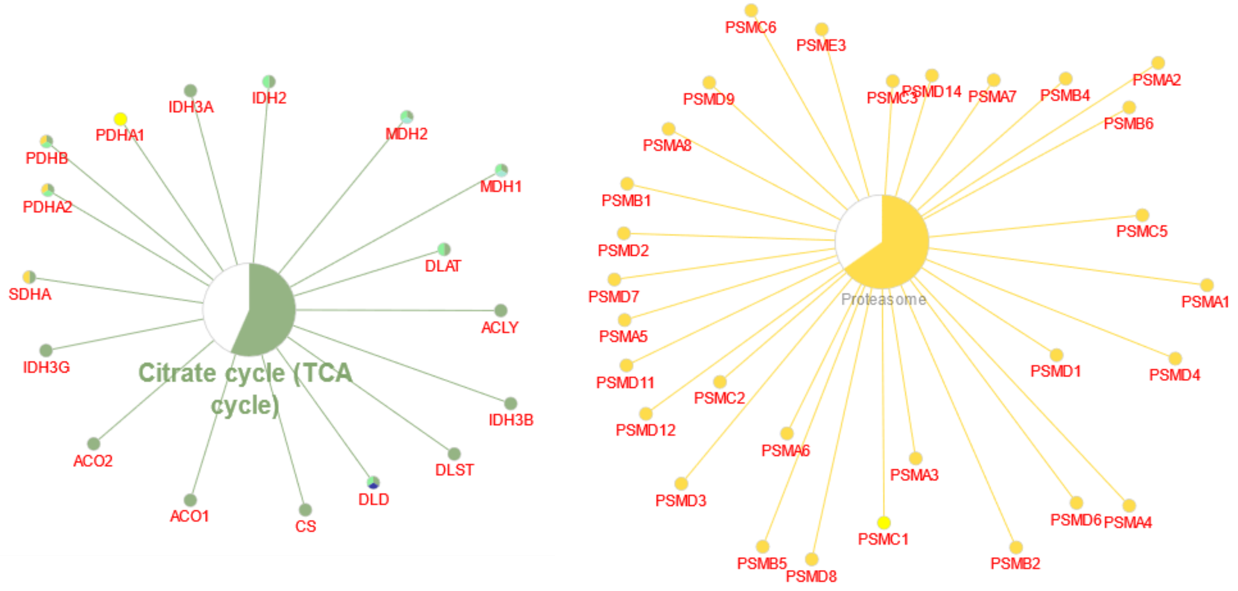


**Figure S2. Enriched KEGG pathways, including all protein members identified by SMAD of two specific pathways, with more than 50% coverage (matching Fig. 2c).** Pathway enrichment analysis was done in Cytoscape with the plugin clueGO. Colored portion of the circle gives the proportion of proteins in that pathway that were identified.

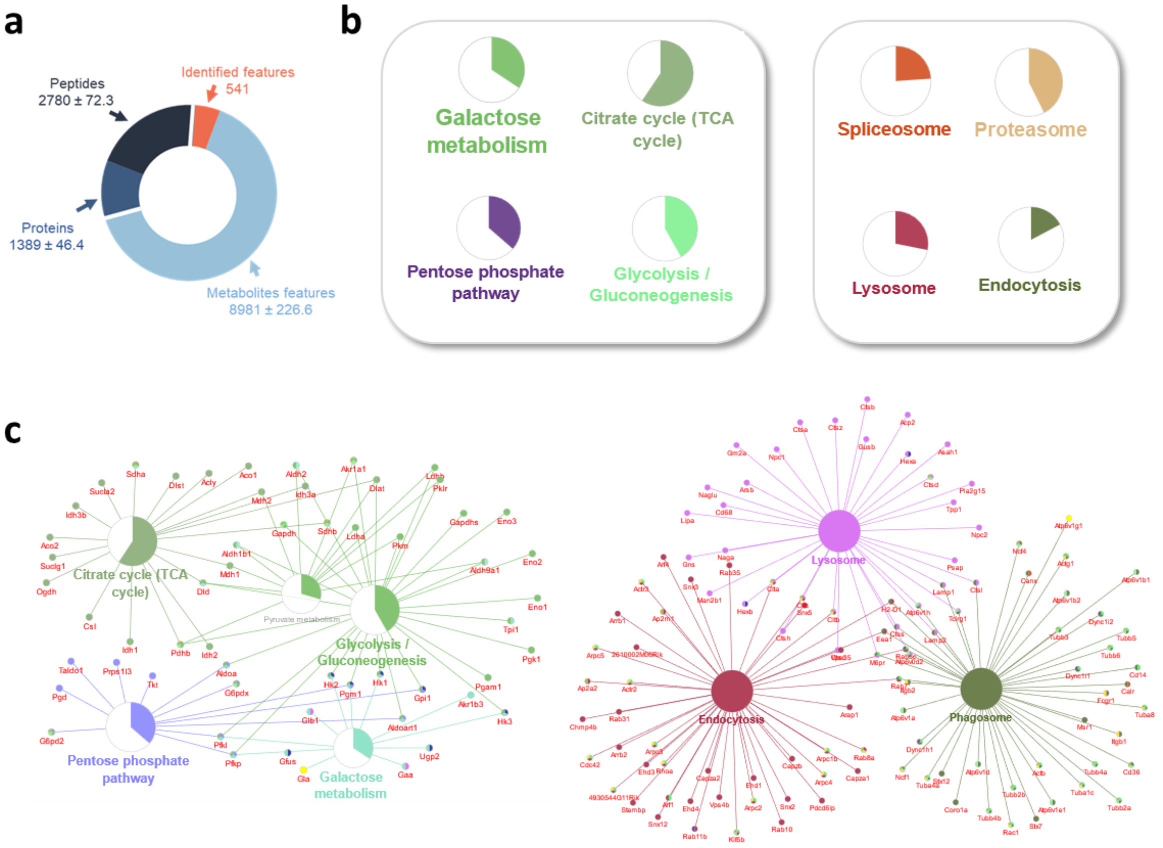


**Figure S3. Performance of SMAD in macrophages. a,** Number of detected metabolite features, peptides, and proteins of macrophages by SMAD. **b,** KEGG pathway enrichment analysis of the proteins identified from macrophages by SMAD. The bars indicate how many proteins in the pathway were identified, and the colored proportion of the circle reflects the coverage of the proteins in each pathway. **c,** Proteins related to typical KEGG pathways identified by SMAD from macrophages.

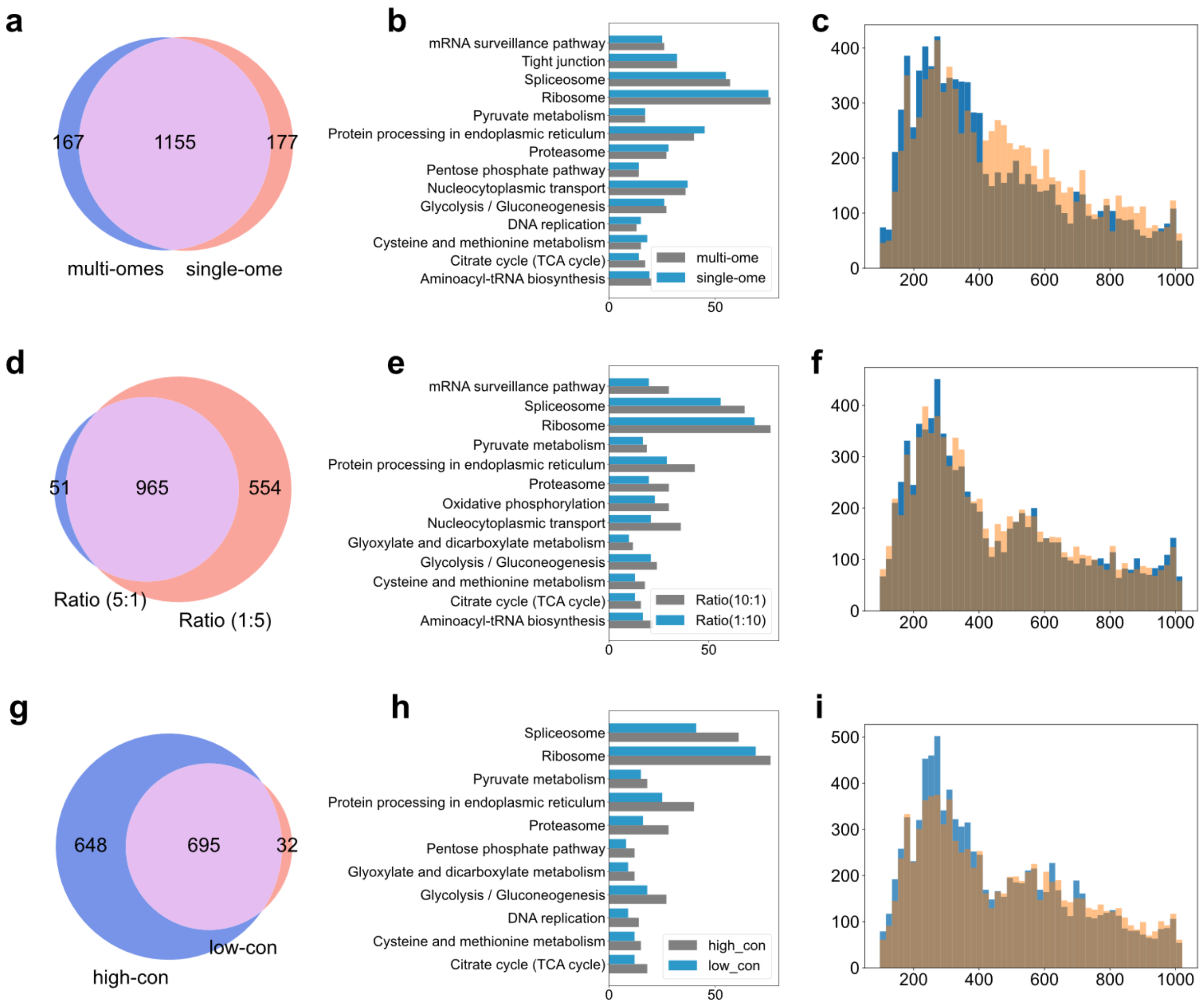


**Fig. S4|Comparison of protein species, enriched pathways, and m/z distribution between different conditions.** **a,** Venn diagram showing the overlap of protein species between the multi-omics method and single-omic methods. **b,** KEGG enriched pathways showing that basically no difference between multi-omics method and single-omic method. **c,** comparison of m/z distribution of detected metabolite features between multi-omics method and single-omic method. **d,** Venn diagram showing the overlap of protein species between high metabolome/proteome ratio (5:1) and low metabolome/proteome ratio (1:5). **e,** KEGG enriched pathways showing that the differences between high metabolome/proteome ratio (:1) and low metabolome/proteome ratio (1:5). **f,** comparison of m/z distribution of detected metabolite features between high metabolome/proteome ratio (5:1) and low metabolome/proteome ratio (1:5). **g,** Venn diagram showing the overlap of protein species between high sample concentration and low sample concentration. **h,** KEGG enriched pathways showing the differences between high sample concentration and low sample concentration. **i,** comparison of m/z distribution of detected metabolite features between high sample concentration and low sample concentration.

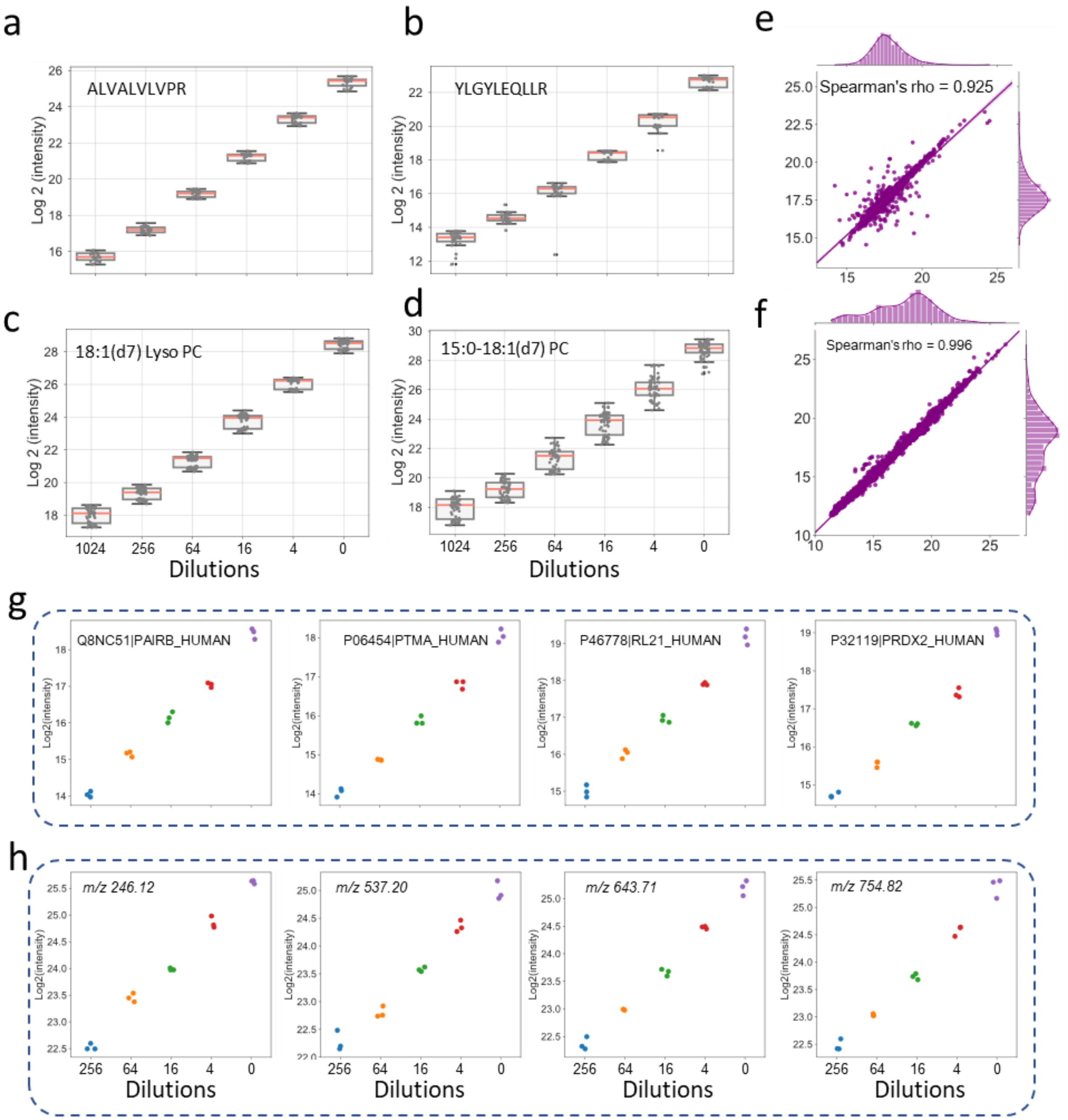


**Fig. S5| Quantification assessment of SMAD with standards and real samples. a,b,** label free quantification curve of two standard peptides from MS-QCAL protein spiked in real samples. (each ratio was measured in triplicate). **c,d,** label free quantification curve of two standard lipids from Avanti spiked in real samples. (each ratio was measured in triplicate). **e,f,** Scatterplot of peptide (e) and metabolite feature intensities (f) quantified by SMAD from two injections of multi-omic samples of 293T cells. **g,** examples of typical quantified proteins from real samples. h, examples of typical quantified metabolite features from real samples.

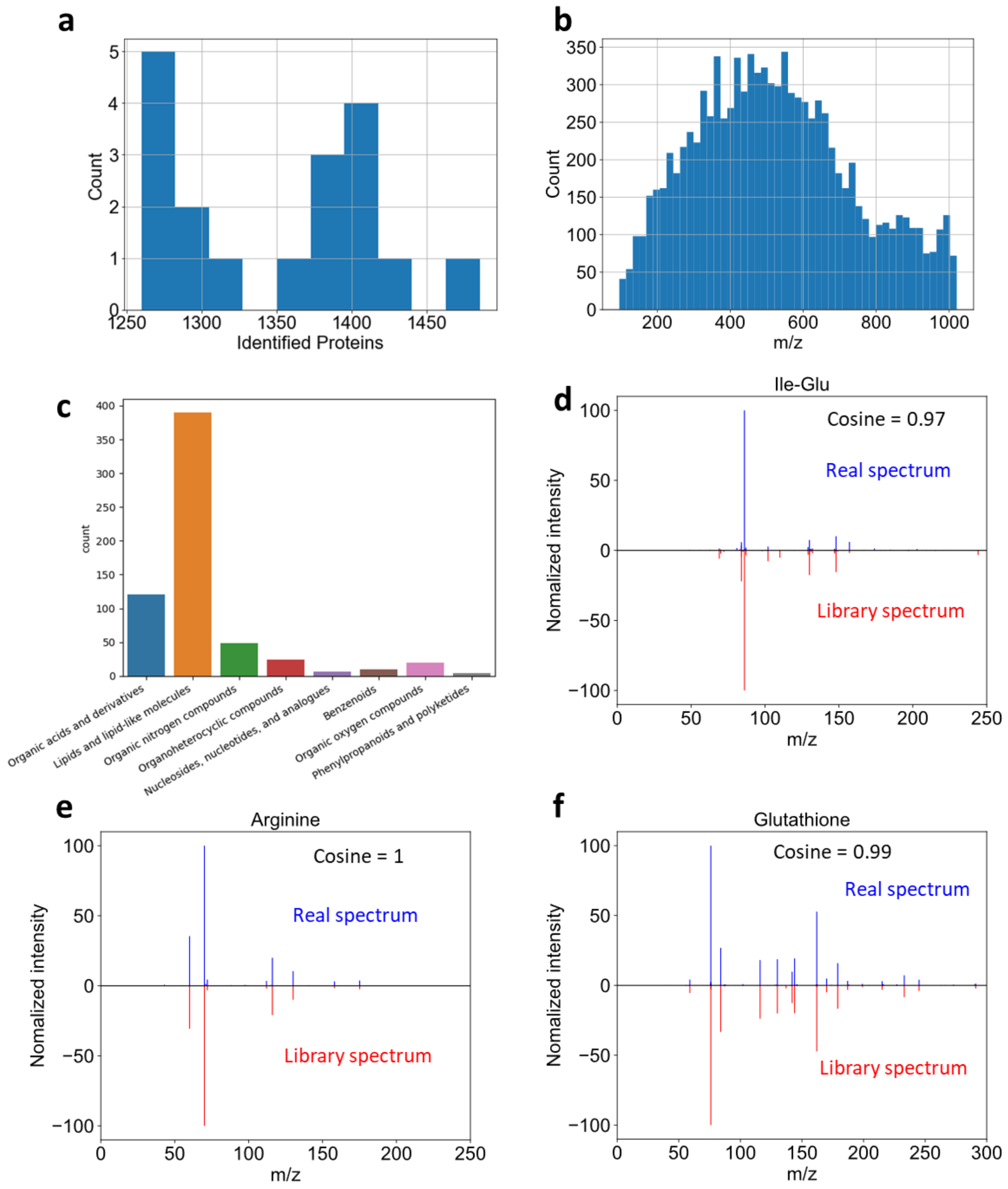


**Figure S6. Macrophage dataset evaluation and metabolite identifications. a,** Histogram of identified proteins in each injection, including all treatments and replicates. **b,** Histogram of m/z distribution of all detected m/z features in macrophages. **c,** class of identified metabolite features in macrophages. **d,e,f,** Tandem mass spectra matching plot for typical metabolites including Ile-Glu(d), Arginine(e) and Glutathione(f).

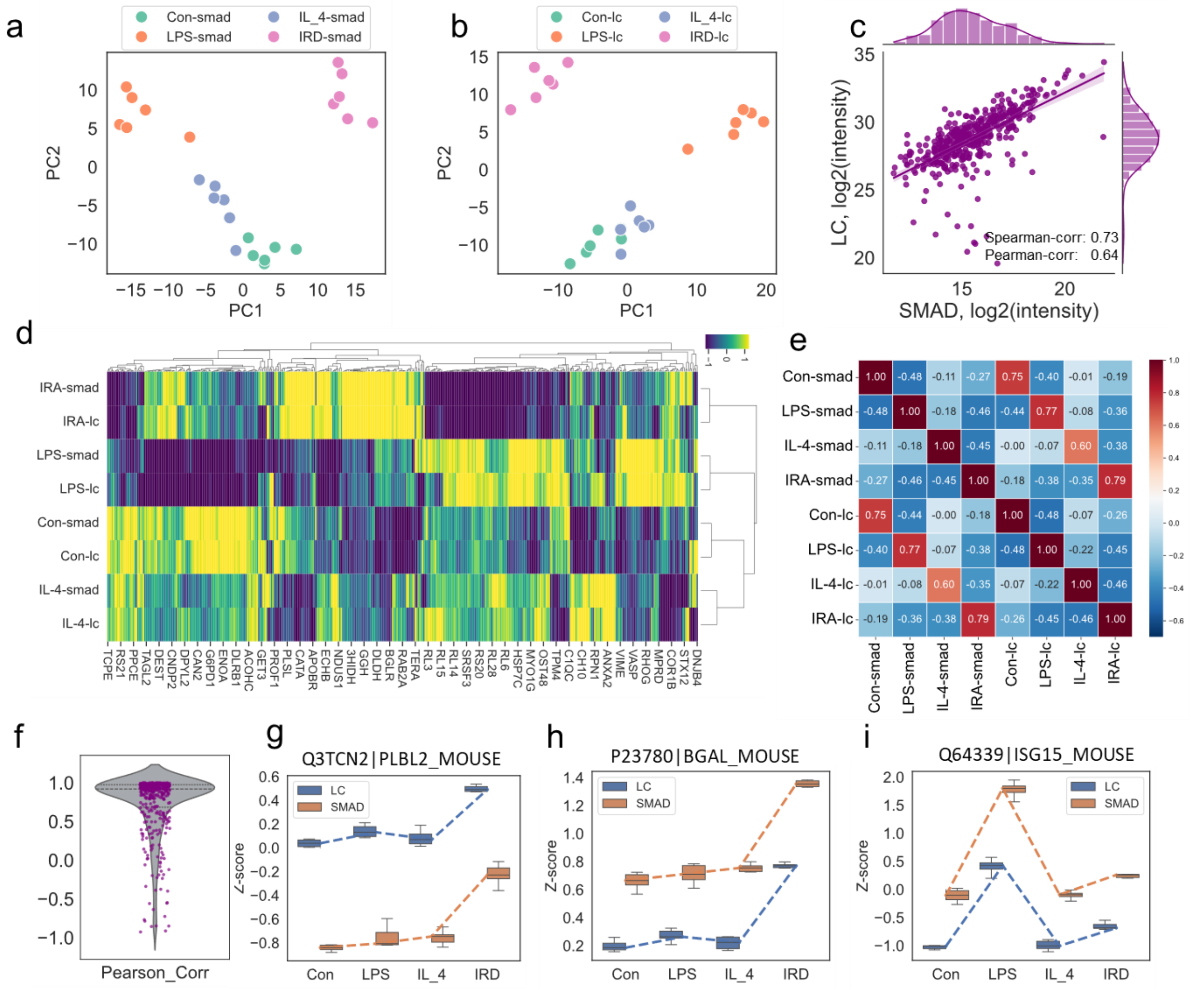


**Fig. S7| Comparison of proteomics quantification results between the SMAD method and the traditional LC method on macrophages after different polarization.** (**a, b**) PCA analysis demonstrates that the datasets generated by the two methods exhibit consistent distribution patterns across different treatment groups. (**c**) Quantitative correlation analysis of proteins (log2 transformed intensity) identified by both methods within the same treatment group is presented (Control group as an example). (**d**) Clustering heatmap analysis of co-identified proteins reveals similar patterns within treatments and consistent trends in their changes between treatments across methods. (**e**) Pearson correlation analysis highlights the relationships between different treatment groups and data acquisition methods (proteins were Z-scored across treatments). (**f**) The distribution of the Pearson correlation coefficients indicates that the identified proteins' change patterns under different treatments are essentially consistent between the two methods. (**g,h,i**) Representative examples of three proteins illustrate consistent quantitative trends between the two methods.

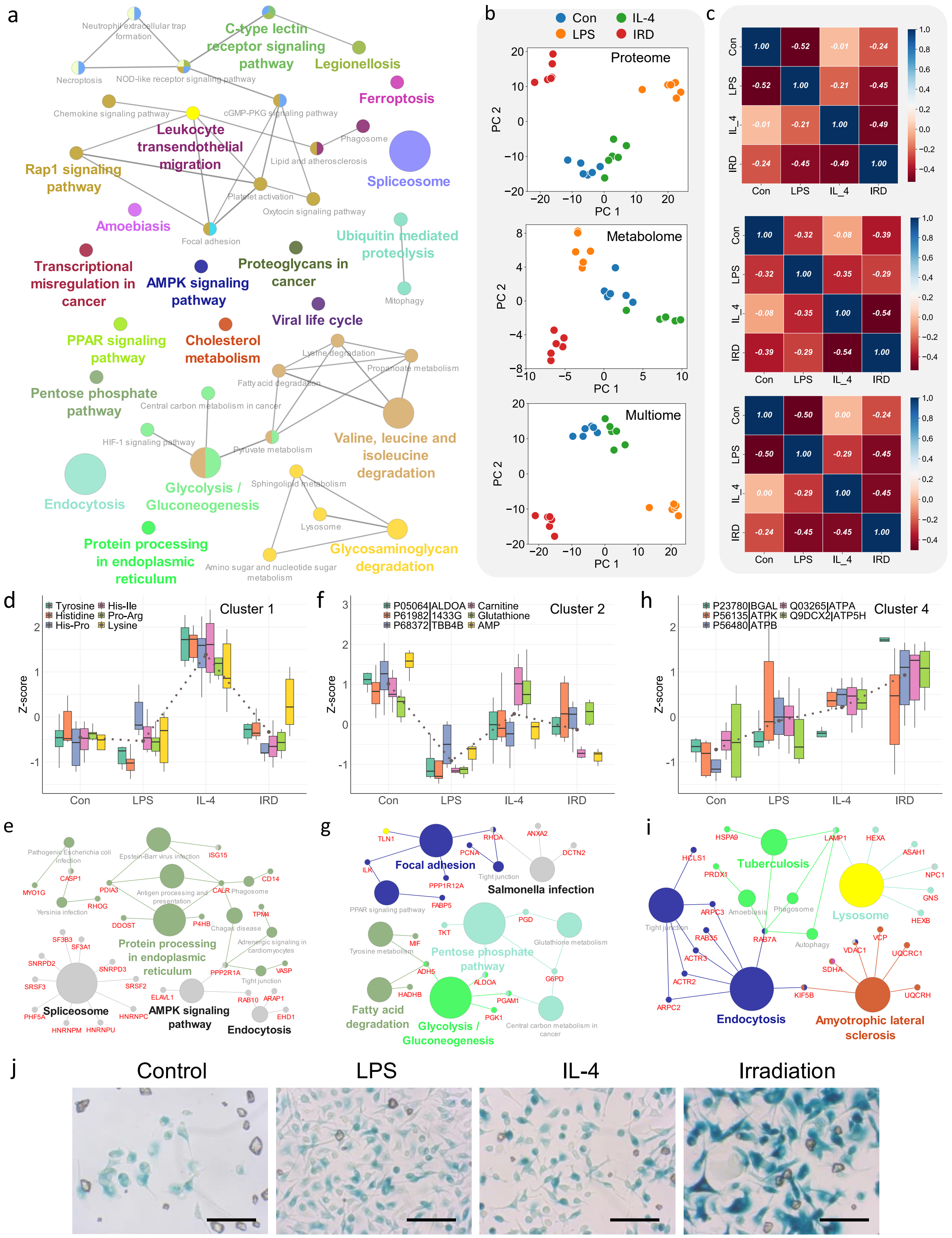

a
C-type lectin receptor signaling pathway
Legionellosis
Ferroptosis
Leukocyte transendothelial migration
Spliceosome
Rap1 signaling pathway
Amoebiasis
Ubiquitin mediated proteolysis
Transcriptional misregulation in cancer
AMPK signaling pathway
Proteoglycans in cancer
Viral life cycle
PPAR signaling pathway
Cholesterol metabolism
Pentose phosphate pathway
Valine, leucine and isoleucine degradation
Glycolysis / Gluconeogenesis
Endocytosis
Protein processing in endoplasmic reticulum
Glycosaminoglycan degradation
b
Con
LPS
IL-4
IRD
Proteome
Metabolome
Multiome
PC 1
PC 2
c
Con
LPS
IL_4
IRD
d
Cluster 1
Tyrosine
Histidine
His-Pro
His-Ile
Pro-Arg
Lysine
Z-score
f
Cluster 2
P05064|ALDOA
P61982|1433G
P68372|TBB4B
Carnitine
Glutathione
AMP
h
Cluster 4
P23780|BGAL
P56135|ATPK
P56480|ATPB
Q03265|ATPA
Q9DCX2|ATP5H
e
Spliceosome
AMPK signaling pathway
Endocytosis
Protein processing in endoplasmic reticulum
g
Focal adhesion
Salmonella infection
Pentose phosphate pathway
Fatty acid degradation
Glycolysis / Gluconeogenesis
i
Tuberculosis
Lysosome
Endocytosis
Amyotrophic lateral sclerosis
j
Control
LPS
IL-4
Irradiation

**Fig. S8|** (a) KEGG pathway enrichment analysis of significantly changed proteins after different polarization or irradiation in Macrophages. (b) PCA dimensionality reduction reveals the distribution patterns of macrophages under different treatments at the proteomic, metabolomic, and multi-omics levels. (c) Correlation of macrophages under different treatments at the proteomic, metabolomic, and multi-omics levels. (d) Representative dysregulated molecules in cluster 1. (e) Typical enriched KEGG pathways in cluster 3 of case study1. (f) Representative dysregulated molecules in cluster 2. (g) Typical enriched KEGG pathways in cluster 3 of case study1. (h) Representative dysregulated molecules in cluster 4. (i) Typical enriched KEGG pathways in cluster 4 of case study1. (j) Enzymatic colorimetric reaction images demonstrate a marked increase in β-galactosidase activity following irradiation treatment.

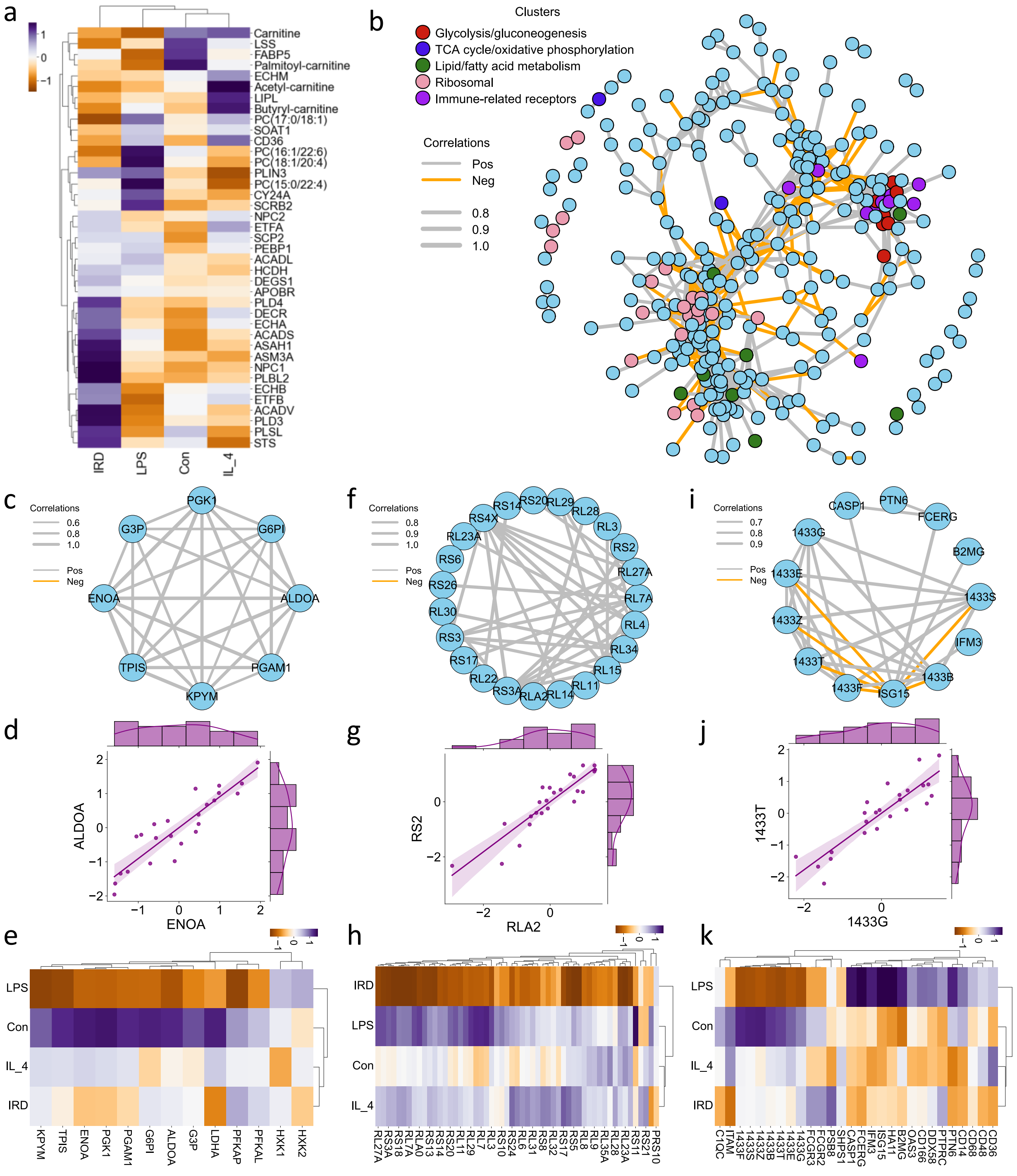

**Fig. S9**| (**a**) Heatmap illustrating the dysregulation of molecules across multi-omic layers involved in lipid metabolism. (**b**) Network representation of all identified molecules with strong correlations (edges filtered by >0.8); molecules associated with the same pathway tend to cluster together. (**c**) Subnetwork of glycolytic proteins. (**d**) Pairwise correlation plot of selected glycolytic proteins. (**e**) Heatmap depicting the dysregulation patterns of all glycolysis-related proteins. (**f**) Subnetwork of ribosomal proteins. (**g**) Pairwise correlation plot of selected ribosomal proteins. (**h**) Heatmap showing the dysregulation patterns of all identified ribosome-related proteins. (**i**) Subnetwork of immune-related receptors and signaling proteins. (**j**) Pairwise correlation plot of selected immune-related and signaling proteins. (**k**) Heatmap illustrating the dysregulation patterns of all identified immune-related and signaling proteins.

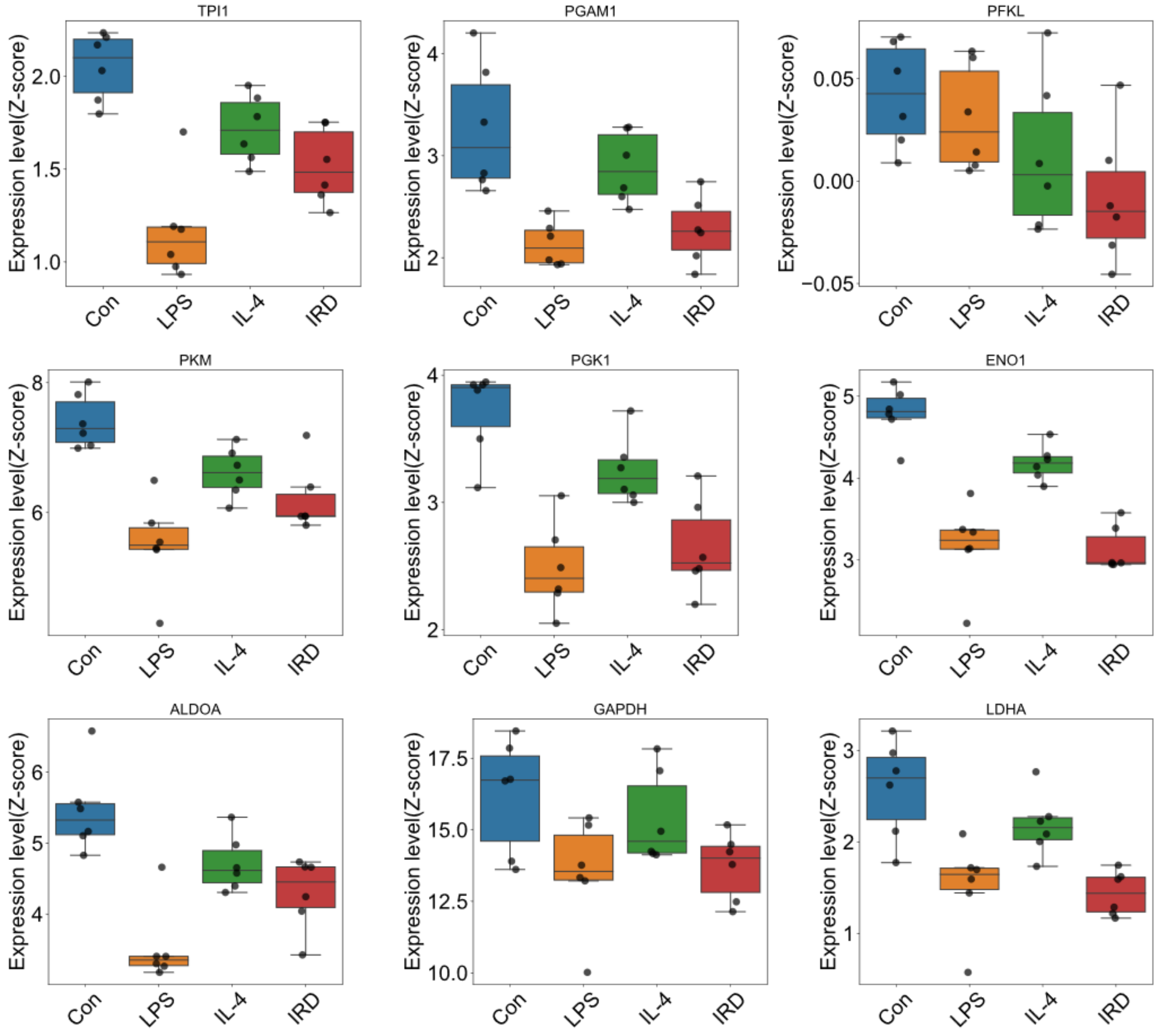


**Figure S10. Dysregulation pattern of glycolytic related proteins identified by LC-MS results.** 9 proteins related to glycolytic validated by LC-MS results (n=6).

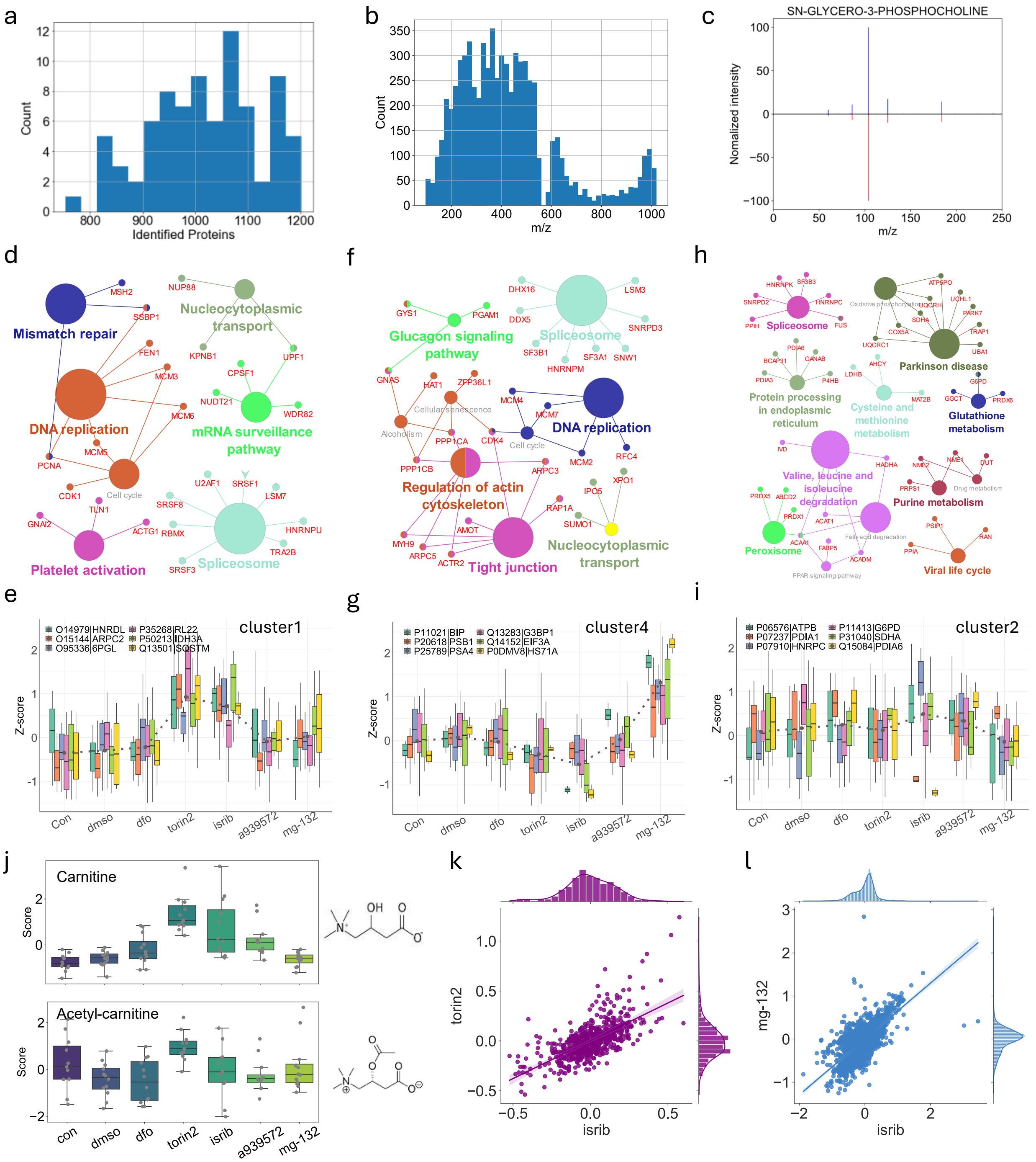


**Fig. S11| Data quality, Pathway analysis for clusters in case study 2. a,** Histogram of identified proteins in each injection including all drug treatments and replicates. **b,** Histogram of m/z distribution of all detected m/z features in case study2. **c,** Tandem mass spectra matching plot for typical metabolites SN-GLYCERO-3-PHOSPHOCHOLINE. **d,** Typical enriched KEGG pathways in cluster 1 of case study2. **e,** Representative dysregulated molecules in cluster 1. **f,** Typical enriched KEGG pathways in cluster 4 of case study2. **g,** Representative dysregulated molecules in cluster 4. **h,** Typical enriched KEGG pathways in cluster 2 of case study2. **i,** Representative dysregulated molecules in cluster 2. **j,** Boxplot of Carnitine(top) and

Acetyl-Carnitine(bottom) in all drug treatments and controls. **k,** Scatterplot of correlation between drug pairs (Isrib and Torin2) from proteome data. **l,** Scatterplot of correlation between drug pairs (MG-132 and Isrib) from metabolome data.

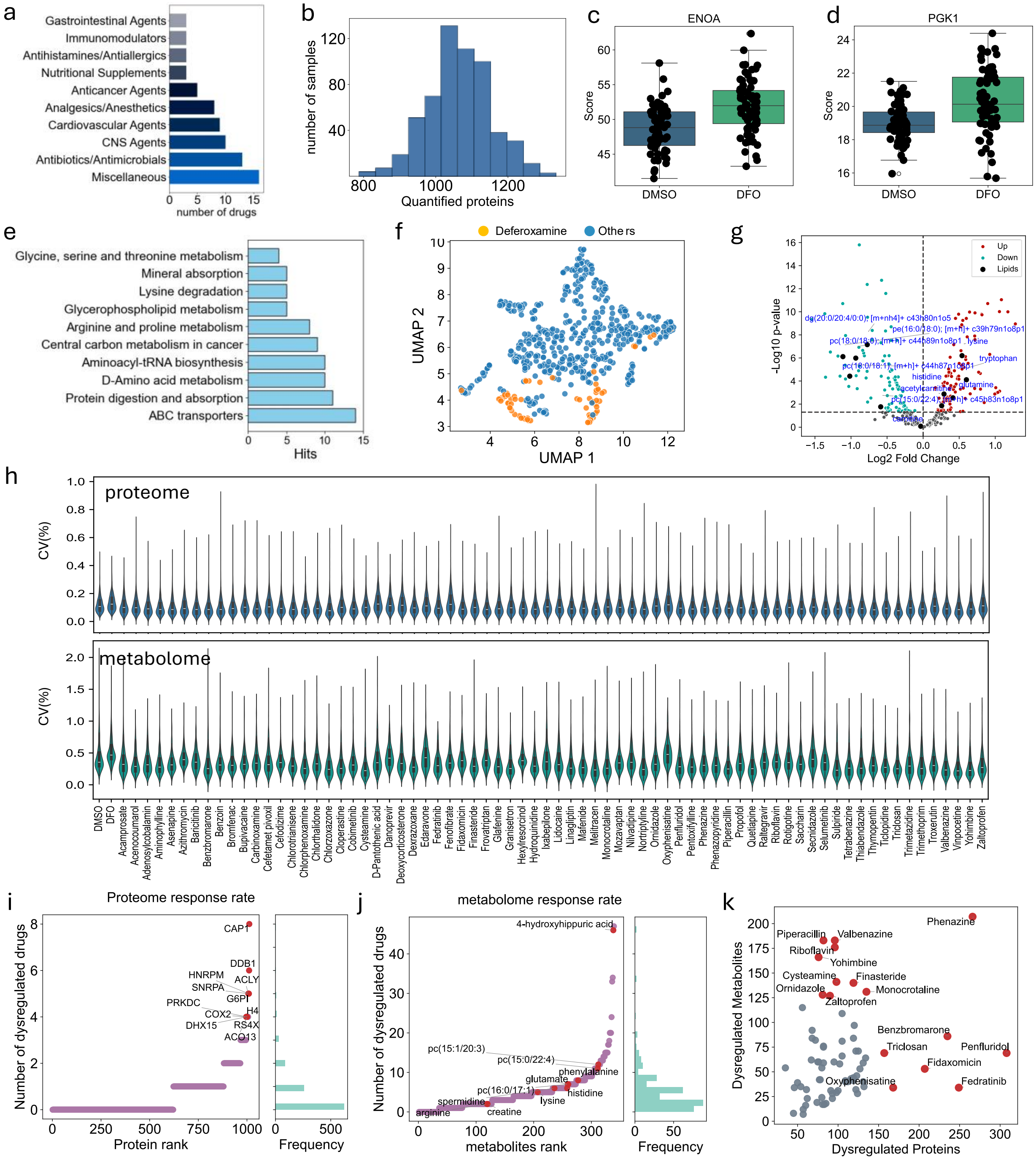


**Extended Data Fig. S12| Data quality and multi-omics response of SMAD in large scale drug screening experiment. a,** Classification of compounds used in the drug screening experiment. **b,** Histogram of quantified proteins in each sample including all drug treatments and replicates. **c,d,** Box-scatter plot of two typical glycolytic proteins that significantly dysregulated in negative control(DFO treatment). **e,** Top enriched pathways of identified metabolites by SMAD in case study3. **f,** UMAP analysis of negative control drug DFO for metabolome result. **g,** Volcano plot of dysregulated metabolites after DFO treatment. **h,** Coefficient of variance distribution for all identified proteins and metabolite features across all replicates within each treatment. **i,** Perturbation-based ranking of all identified protein features across different drug treatments. **j,** Perturbation-based ranking of all identified metabolite features across different drug treatments. **k,** Dysregulated proteins and metabolites for each drug without Benjamini–Hochberg (BH) adjusted P values <0.05.

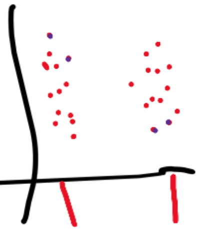

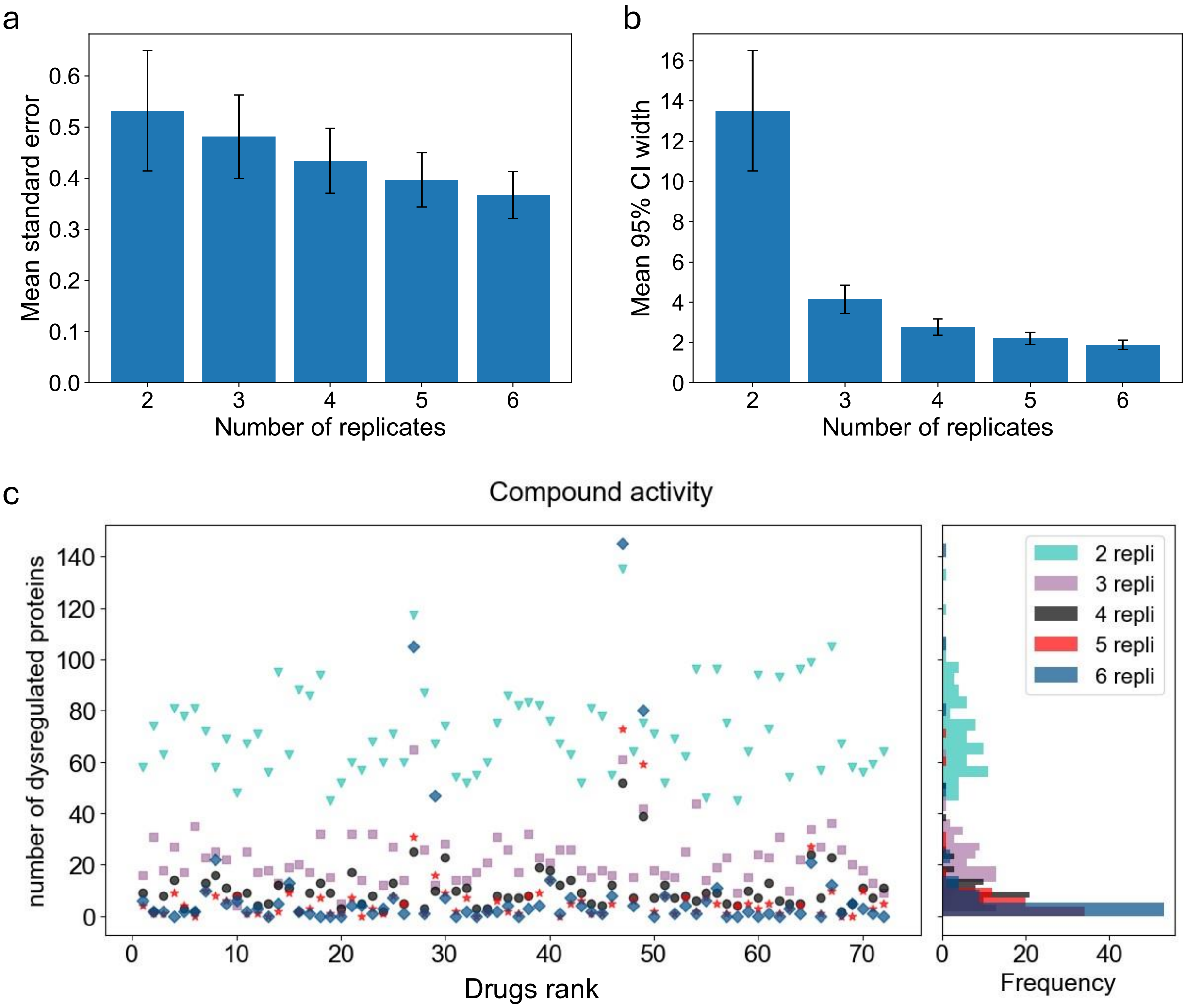


**Figure S13. Influence of replicate number on data quality and dysregulation profiling of drug screening dataset. a,** Bar plot showing the average standard error (± standard deviation) across all proteins as a function of the number of biological replicates (n). Each value is derived from 200 bootstrap samples across all 72 treatment groups. Increasing the number of replicates leads to a monotonic decrease in SE, indicating improved precision in protein quantification. **b,** Bar plot showing the average width of the 95% confidence intervals (± standard deviation) for protein abundance estimates, computed from 200 bootstrap replicates for each group size. Confidence intervals become narrower as N increases, demonstrating that higher biological replication yields more reliable and constrained estimates. **c,** The average number of significantly dysregulated proteins (Significance calculated by student T-test with Benjamini–Hochberg (BH) adjusted P values <0.05) identified in pairwise comparisons between each drug treatment and control, as a function of the number of biological replicates (n).

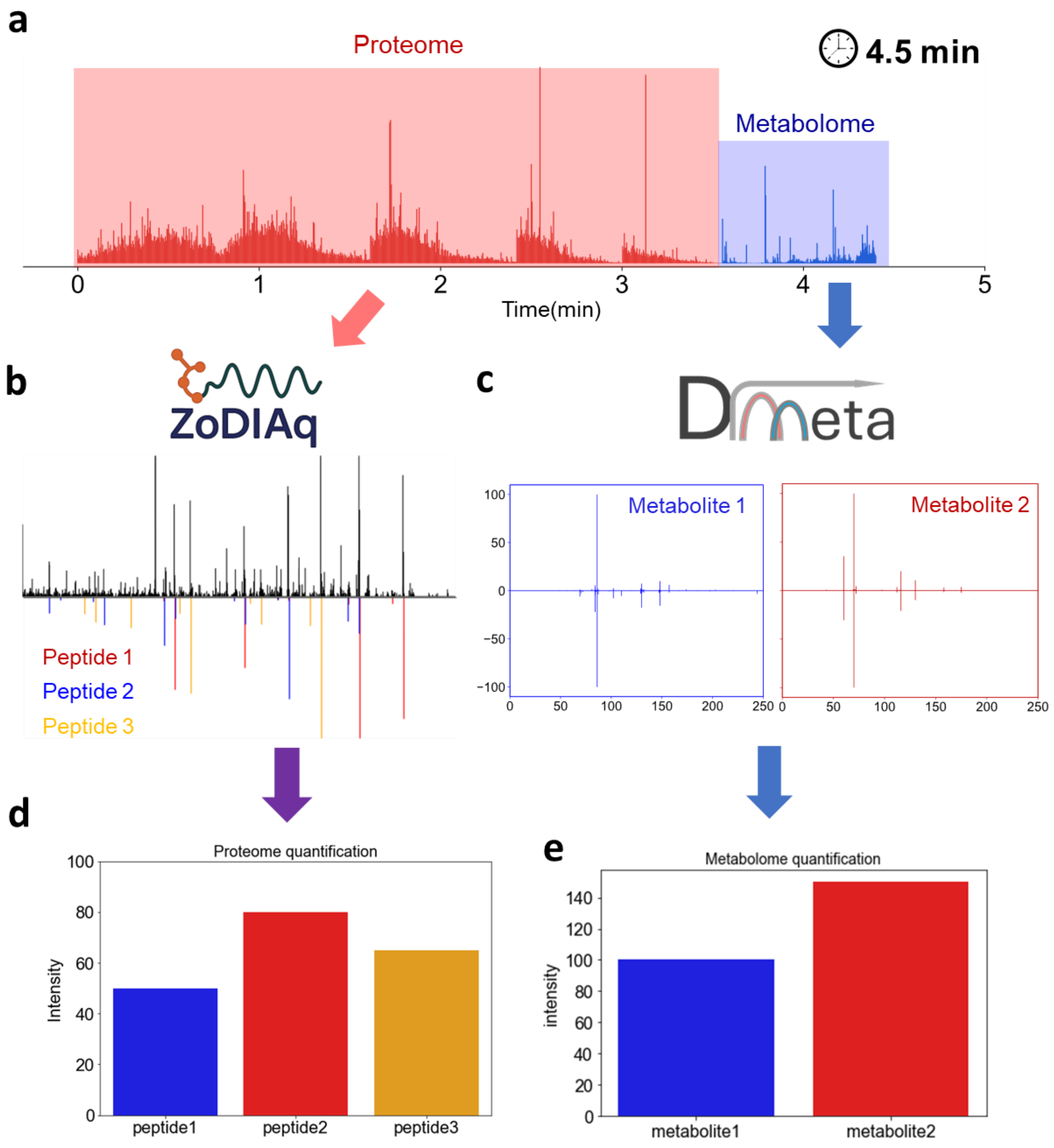


**Figure S14. Quantitative and identification schematic diagram of SMAD in case study 3. a,** Typical TIC figure of SMAD. **b,** identification of peptides from tandem mass spectra by spectra-to-spectra match. **c,** identification of metabolites from tandem mass spectra by spectra-to-spectra match. **d,** Quantification of proteome by summing up all fragment intensities of a specific peptide. **e,** Quantification of metabolome by summing up all fragment intensities of a specific metabolite feature.

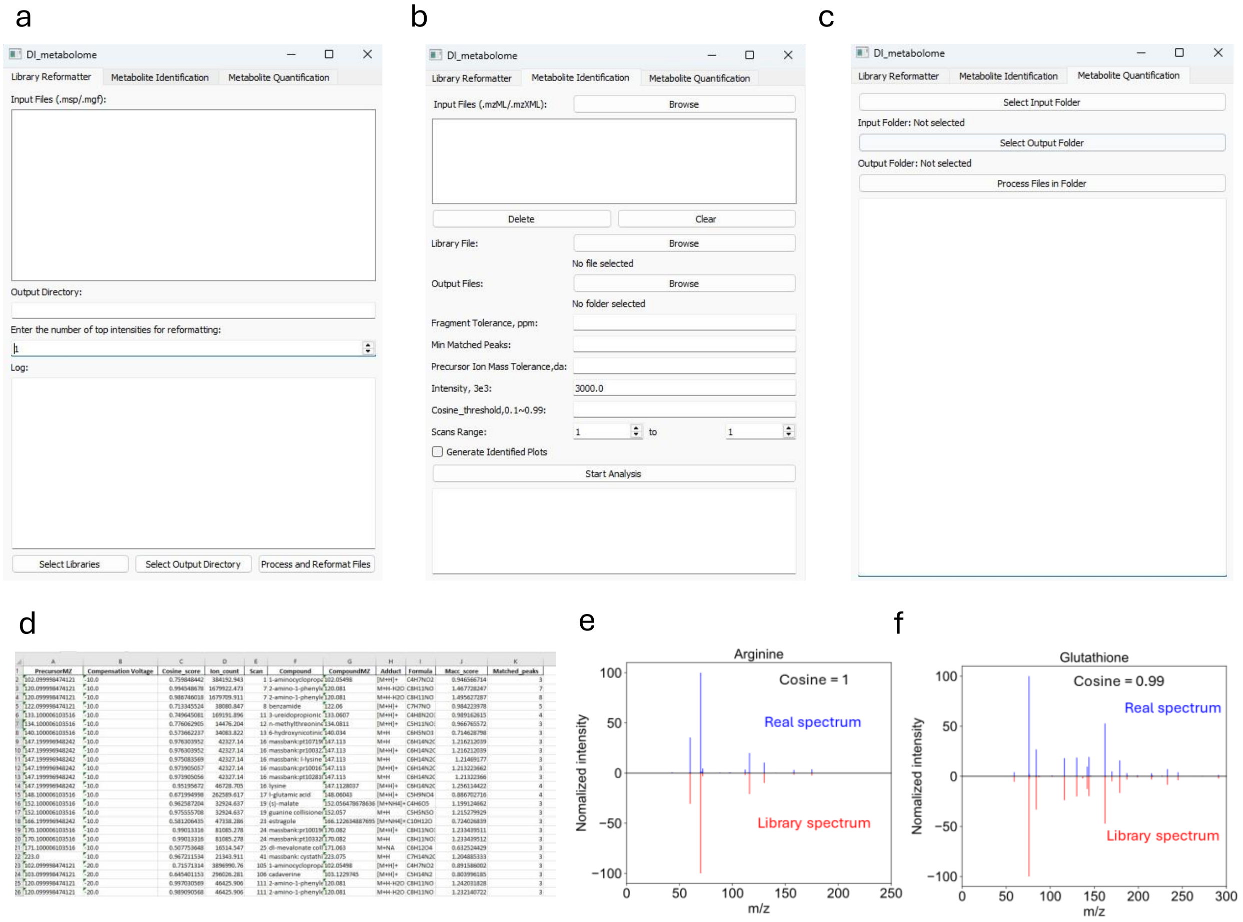


**Extended Data Fig. S15| Demonstratin of DImeta software. a,b,c,** Three modules and corresponding GUI interface of DImeta software designed for metabolites identification of direct injection. Library reformat(a), metabolite identification(b) and Metabolite quantification(c). **d,** The output file format of identification results from DImeta, including precursorMZ, compensation voltage, cosine_score, ion_count, scan_number, compound_name, compoundMZ, adduct type, compound_formula, matched_peaks and macc_score. **e,f,** Tandem mass spectra matching plot for typical identified metabolites from DImeta. Arginine (e) and Glutathione(f).

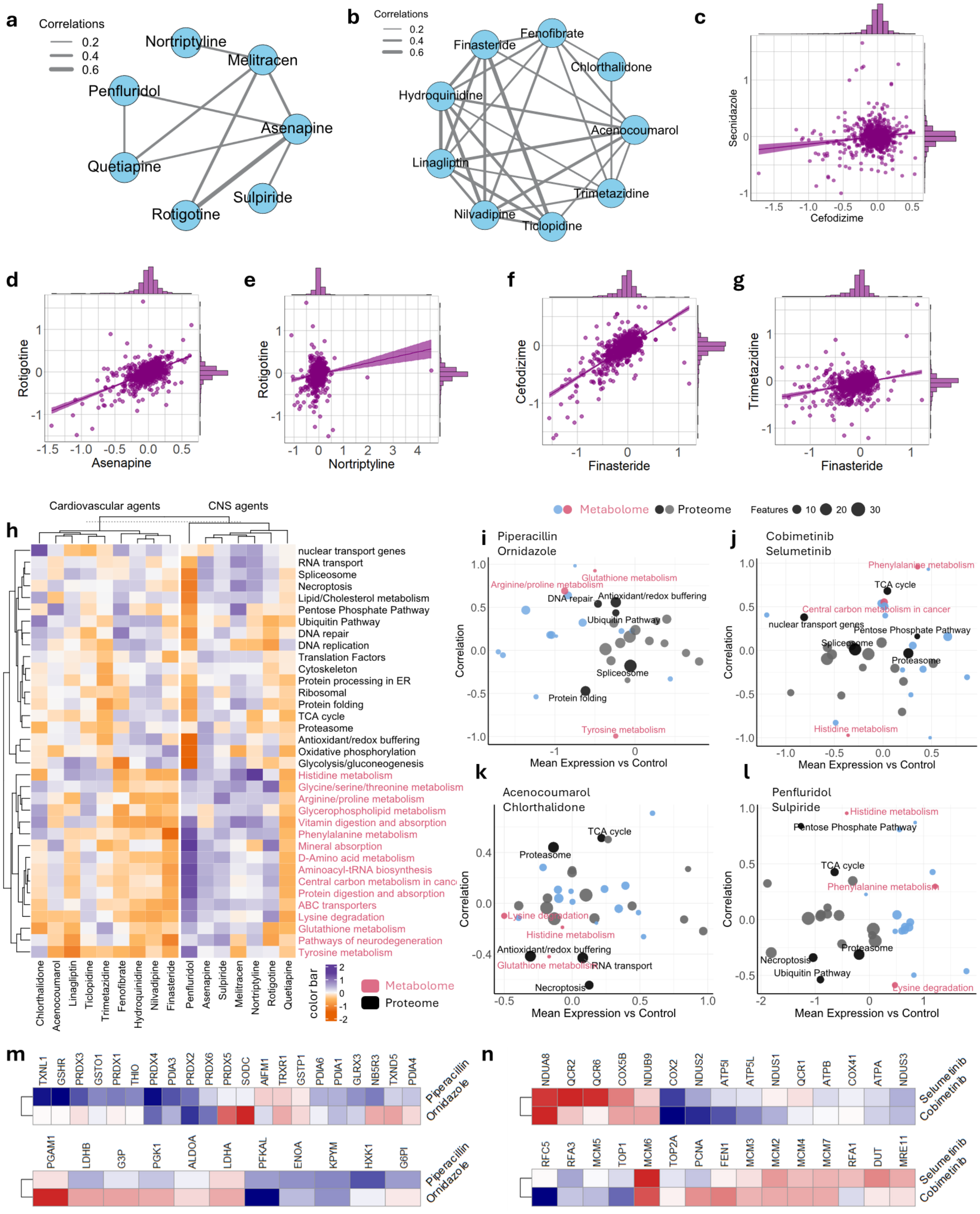


**Fig. S16| a,b,** Subcommunity of central nervous system (CNS) agents(a) and Cardiovascular agents(b). **c,** Pairwise correlation plot of two antibiotics secnidazole and cefodizime showing example of relatively weak correlation. **d,e,** Pairwise correlation plot of typical compounds in CNS cluster demonstrating moderate(d) and weak correlation(e). **f,g,** Pairwise correlation plot of typical compounds in cardiavacular cluster demonstrating moderate(f) and weak correlation(g). **h,** Clustered heatmap illustrates the changes in multi-omics pathways across different drug clusters (CNS agents and Cardiovascular agents), with metabolomics (red) and proteomics (black). **i,j,k,l,** Correlation and perturbation patterns of antibiotics(i), MEK inhibitors(j), CNS agents(k), and Cardiovascular agents(l) across proteomic and metabolomic pathways. **m,n,** changes in associated molecules within representative dysregulated pathway related to two antibiotics(m) and two MEK inhibitors(n).

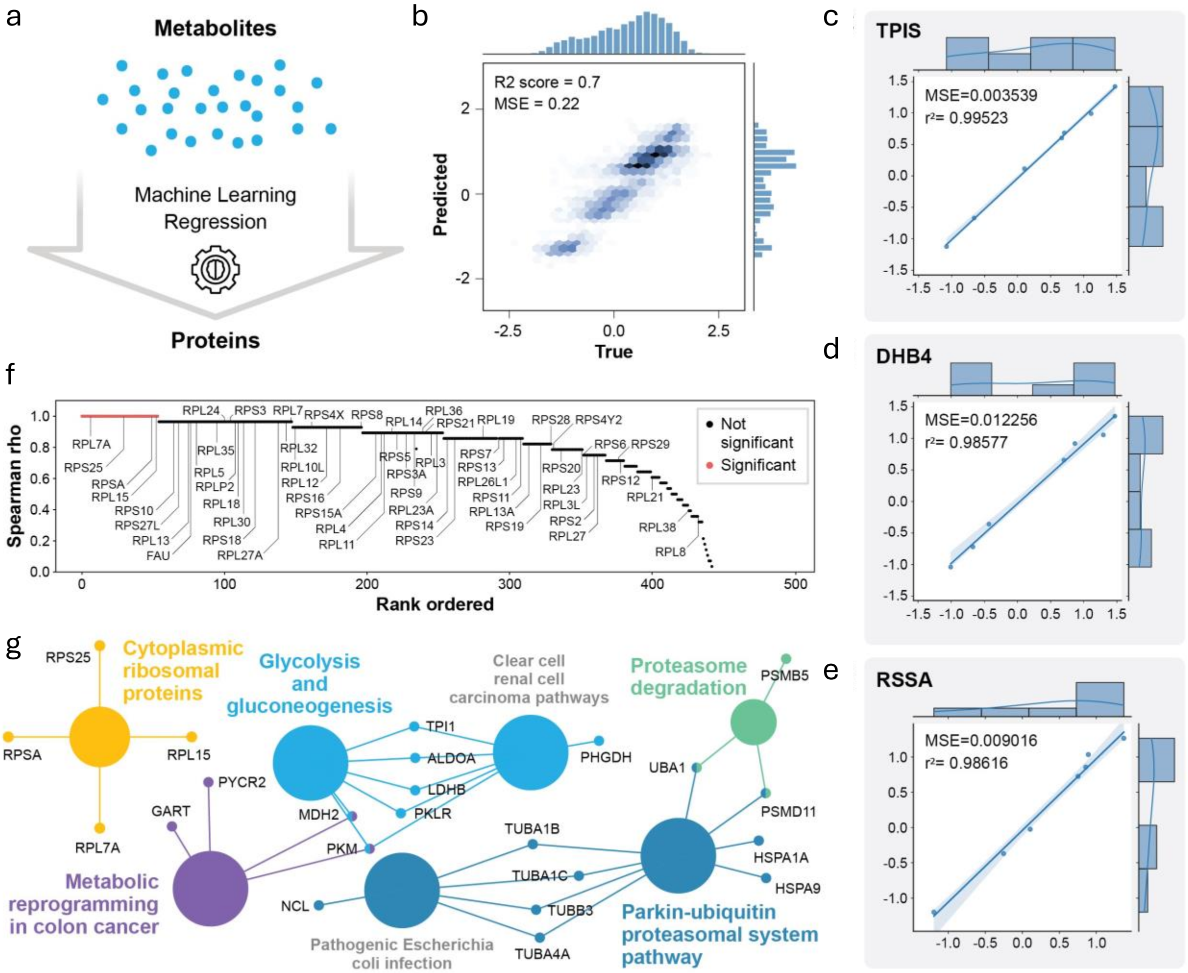

**Fig. S17| Machine learning predicts proteomic changes from metabolomic changes. a**, Scheme showing the ML task of predicting proteins from metabolomic data measured in parallel by SMAD. **b**, Overall performance of all proteins true versus ML predicted values. **c,d,e**, Examples of three of the most well predicted proteins from different pathways: TPIS from glycolysis, DHB4 from fatty acid oxidation, and RSSA from the ribosome. **f**, plot of the Spearman rho between true versus predicted protein quantities for all 450 proteins. Red indicates statistically significantly predicted proteins according to a Bonferroni corrected p-value from Spearman correlation analysis. **g**, WikiPathways term enrichment analysis of the 54 significant proteins from (f).

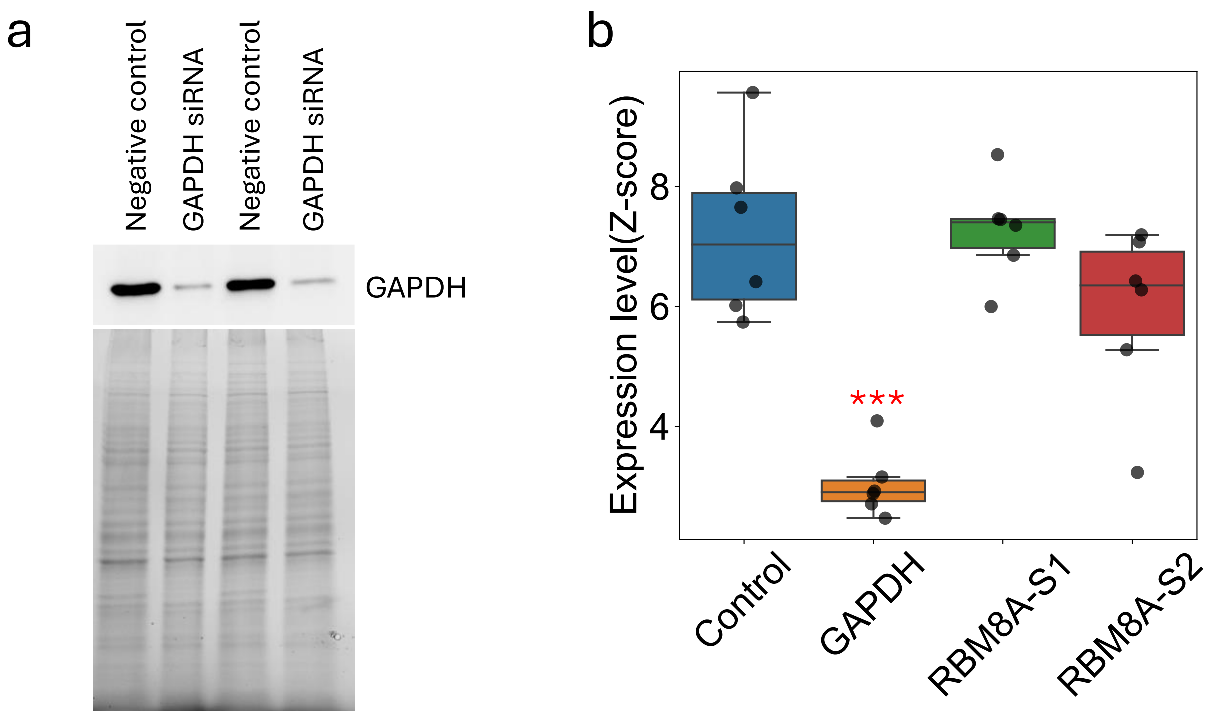


**Figure S18. Validation of GAPDH knockout by Western blot and proteomics quantification.**

**a,** Representative Western blot showing effective knockdown of GAPDH protein expression. GAPDH signal is strongly reduced in knockdown samples compared to a siRNA negative control, confirming successful gene disruption (n=2, biological replicate). The lower panel indicates the total amount of protein per line (TGX Stain-Free Gel, Bio-Rad). **b,** Quantitative plot illustrating GAPDH protein abundance from LC-MS proteomic analysis. Each point represents an individual biological replicate (n=6). GAPDH levels are significantly downregulated in the knockdown group ($p < 0.001$), consistent with the Western blot result.
